# Phage-assisted continuous evolution of enzymes for noncanonical tyrosine biosynthesis

**DOI:** 10.64898/2026.05.08.723366

**Authors:** James S. Andon, Abhijit Behera, Debashrito Deb, Amy M. Weeks, Andrew R. Buller, Tina Wang

**Author notes:** Correspondence should be addressed to Tina Wang. These authors contributed equally and are listed in alphabetical order (James S. Andon, Abhijit Behera).

## Abstract

Genetic code expansion introduces new-to-nature chemical moieties into ribosomally synthesized proteins. In practice, the scope of functional groups that can be accessed using this method is often limited by noncanonical amino acid (ncAA) availability. Producing ncAAs directly in cells can circumvent poor ncAA uptake or commercial unavailability, but limited enzymes suitable for this application exist. *In vitro* evolution campaigns have been remarkably successful in yielding synthetically useful “ncAA synthases.” However, these enzymes are optimized for preparative-scale synthesis and their activities often do not translate well to cellular biosynthesis. Thus, expanding strategies to engineer enzymes specifically for ncAA production within cells will benefit further implementation of genetic code expansion. Here, we use phage-assisted noncontinuous and continuous evolution to evolve enzymes for improved synthesis of non-canonical tyrosine derivatives in *E. coli*. Using simple serial passaging, we uncovered mutations that doubled the production of an expensive ncAA, 3-methoxytyrosine, by tyrosine phenol lyase, and furthermore evolved variants that enable 3-iodotyrosine biosynthesis, a transformation the parent enzyme is unable to catalyze. Additionally, we evolved a recently reported tyrosine synthase for improved production of 3-halogenated tyrosines, identifying variants that exhibit high activity even at low substrate concentrations owing to a ∼8-fold reduction in K_M_. Our results demonstrate that phage assisted evolution can be used to rapidly improve the activity of enzymes for ncAA production in cells.

## Introduction

Proteins produced by nature are generally limited to 20 canonical amino acids, restricting the chemical diversity and functionalities available. Genetic code expansion allows the insertion of noncanonical amino acids (ncAAs) with custom chemical moieties into cellularly synthesized peptides and proteins.^1^ The most widely used methods employ orthogonal aminoacyl-tRNA synthetase (aaRS)/tRNA pairs that enable ncAA installation into a recoded codon (often UAG). These approaches have enabled site-specific incorporation of probes, bioorthogonal handles, and post-translational modifications into target proteins of interest, as well as the construction of artificial metal centers and new-to-nature catalytic sites in designed enzymes, among many other applications.^2,3^

Although engineered aaRS/tRNA pairs have been reported for over 500 ncAAs,^4^ the high costs of ncAAs and their inefficient transport across the cell membrane present hurdles to the accessibility of this technology.^5^ *In situ* biosynthesis and semi-synthesis, where ncAAs are produced from endogenous metabolites or exogenously added precursors, respectively, have gained attention as potential strategies for mitigating these issues by lowering costs and employing precursors that may be more readily membrane-permeable.^6^ Recent examples have showcased the ability of heterologously expressed enzymes to mediate the biosynthesis or semi-synthesis of specific ncAAs in *E. coli*.^7–11^ However, the appeal of genetic code expansion hinges on introducing an ever-expanding range of new-to-nature functional groups into proteins. *In vitro* directed evolution has seen remarkable success for furnishing enzymes that produce ncAAs under *in vitro* conditions, which typically feature > 10 mM substrate loadings and organic co-solvents.^12–16^ However, enzymes evolved under such conditions are often poorly suited for immediate use in cellular biosynthesis.^17,18^

An appealing strategy to evolve ncAA synthases is to use a biosensor that detects ncAAs by way of their incorporation into proteins. By selecting for ncAA incorporation into amber-suppressed antibiotic resistance markers, researchers have identified improved hydroxylases for 5-hydroxytryptophan production and improved carbamoylases to liberate 3-iodotyrosine and 3-nitrotyrosine from their respective *N-*carbamoylated precursors.^19–21^ However, these studies examined enzymes that perform functional group modifications, which have limited ability to generate a broad range of ncAAs. Additionally, although easily adoptable, the directed evolution approaches used rely on manually generated genetic diversity and discrete steps for accomplishing the selection, which limits the number of rounds that can be performed and impedes more extensive evolutions that may be required for challenging targets. Furthermore, typically only one enzyme target and one set of selection conditions can be examined at a time. Finally, spontaneous cellular mutations (for example, mutating the amber stop codon in the antibiotic resistance gene) can generate false positives in these selections which may require additional counter-selection to weed out.

Continuous evolution methods,^22–24^ such as phage-assisted continuous evolution (PACE),^25^ can offer a speedier and more scalable alternative to traditional directed evolution. PACE enables rapid exploration of sequence space with library sizes routinely approaching ∼10^11^ members and, due to its automation of the directed evolution process, enables far more rounds of selection compared to conventional approaches.^26^ A unique advantage of PACE is the separation of the genetic elements encoding the evolving gene of interest (encoded on phage) and the selection (encoded in *E. coli* host cells). Because only the phage are under selection, false positives from mutations to the selection machinery in the host cells do not enrich to any significant extent. However, the application of PACE to small molecule biosynthesis remains underexplored compared to other targets, in part owing to the difficulty of creating selections for the production of small molecules of interest, which must be linked to phage protein pIII production.^27–29^ Ho and colleagues reported the evolution of the pyrrolysine biosynthetic pathway using PANCE, a non-continuous version of PACE, by incorporating pyrrolysine into an amber-suppressed pIII using a pyrrolysine aaRS/tRNA pair.^30^ Recently, Pulschen and coworkers used PACE to evolve the tryptophan halogenase RebH to achieve better catalytic efficiency and dramatically improved expression.^31^ Targeting enzymes capable of achieving more convergent ncAA syntheses, for example through C-C bond forming reactions, could broaden PACE’s applicability to improving the accessibility of ncAA production in cells.

Here, we use phage-assisted evolution to evolve two C-C- bond forming enzymes, tyrosine phenol lyase (TPL) and tyrosine synthase (TyrS),^32^ for the cellular production of tyrosine-derived ncAAs through semi-synthesis using substituted phenols. Our TPL evolutions rapidly furnished enzyme variants with improved activity on a known substrate, 2-methoxyphenol, as well as new activity on 2-iodophenol, on which the parent enzyme shows no reactivity. TyrS proved a more challenging target, which prompted us to explore a variety of strategies for mutagenesis and evolution. While we found that a combination of PANCE and PACE will furnish more active TyrS variants, we also show that combining a pre-generated site-saturation library with continuous mutagenesis can shortcut evolution using the more accessible PANCE selection format. Finally, by tuning selection stringency, we were able to promote the selection of highly active TyrS mutants. Our most evolved variant exhibited broad activation towards 2-halogenated phenols even at very low substrate concentrations (50 µM), owing to large reductions in K_M_.

## Results

### PACE selection for ncAA-synthesizing enzymes

Our strategy for evolving ncAA synthesizing enzymes in PACE is shown in **Figure 1a**. Here, ncAA synthesis by a phage-encoded ncAA synthase is coupled to pIII production through use of an engineered orthogonal aaRS (o-aaRS) that recognizes the ncAA and charges it onto an orthogonal tRNA (o-tRNA) bearing an anticodon loop decoding the amber stop codon (UAG). The charged o-tRNA suppresses amber stop codons in *gIII*, enabling translation of full-length pIII by the ribosome and subsequent production of infectious progeny phage.^30,33^

**Figure 1.**
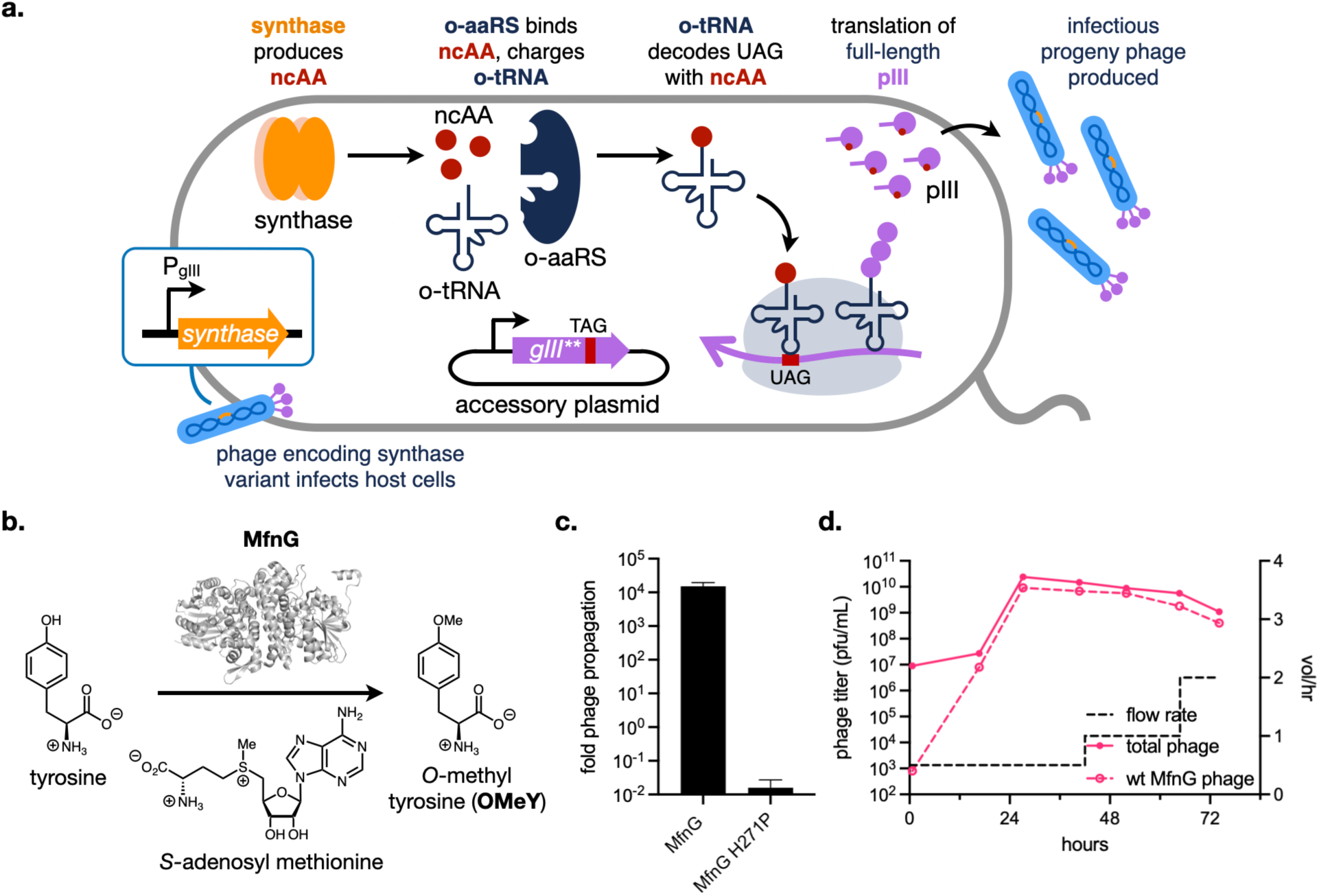
Overview and characterization of a PACE selection for ncAA production. **a.** Overview of the selection strategy. Phage carrying a ncAA synthase in place of *gIII* are used to infect host cells encoding an orthogonal aaRS/orthogonal tRNA (o-aaRS/o-tRNA) pair engineered to charge the target ncAA, and an accessory plasmid (AP) containing *gIII* amber-suppressed at residues P29 and Y184 (*gIII\*\**).^33^ **b.** *Streptomyces drozdowiczii* SCSIO 10141 MfnG catalyzes the SAM-dependent *O*-methylation of tyrosine (PDB ID: 7UX8). **c.** Activity of optimized phage encoding MfnG or an inactivated H271P mutant in 6-hour propagation assays using host cells carrying an AP with *gIII*\*\*. **d.** Phage assisted continuous selection of a 1:10,000 ratio of phage encoding wild-type MfnG to phage encoding the H271P mutant using host cells carrying an AP with *gIII\*\**. Wild-type MfnG phage were quantified by activity-dependent plaquing on host cells.

Our initial tests of this system used the SAM-dependent methyltransferase MfnG, which was recently used to biosynthesize *O*-methyl tyrosine (OMeY) in *E. coli* from tyrosine (**Figure 1b**).^34^ When expressed from plasmid, MfnG robustly biosynthesizes OMeY, which can then be incorporated into amber-suppressed GFP by an engineered *M. janaschii* TyrRS/tRNA pair (*Mj*OMeYRS/*Mj*tRNA^mut^).^35^ We tested if phage encoding the gene for MfnG would be able to trigger pIII production from host cells carrying plasmids bearing *Mj*OMeYRS/*Mj*tRNA^mut^ and *gIII* bearing two amber stop codons (*gIII\*\**). MfnG phage exhibited ∼10,000-fold phage propagation activity on these host cells, indicative of robust ability to produce OMeY resulting in full-length pIII production (**Figure 1c**). When we installed an inactivating mutation (H271P) in MfnG’s active site, phage propagation dropped by > 10^5^-fold (**Figure 1c**), demonstrating absolute dependence of phage fitness on enzyme activity.

Next, we assessed whether our strategy would enable us to select for active MfnG over inactive mutants. We competed phage encoding the inactive H271P MfnG mutant with phage encoding wild-type MfnG in a continuous flow experiment performed without mutagenesis (**Figure 1d**). Despite starting at a 1:10,000 ratio to the H271P mutant, wild-type MfnG phage overtook the population in < 24 hours, as measured by activity-dependent plaque formation on *gIII\*\** host cells. In addition, we attempted to revert the H271P mutation using PACE (**SI Figure 1a**). We observed complete reversion of the proline to histidine in the sequenced phage pools after 24 hours (**SI Figure 1b**). Together, these results suggested to us that this selection strategy could be used to evolve ncAA-synthesizing enzymes through phage-assisted evolution.

### Evolution of tyrosine phenol lyase for increased biosynthesis of tyrosine derivatives

We next examined if we could use our selection strategy to evolve improved variants of tyrosine phenol lyase (TPL), a pyridoxal 5’-phosphate (PLP) dependent C-C bond forming enzyme that is frequently employed to synthesize non-canonical tyrosine derivatives *in vitro* and in cells from phenol, pyruvate, and ammonium (**Figure 2a**).^36–42^ The thermodynamics of the TPL reaction favor amino acid breakdown and is typically only run in the synthetic direction with a high excess of substrate (1-10 mM range from previous studies). Elevated phenol concentrations are toxic to cells and undesirable for biosynthesis. We therefore asked whether TPL could be evolved to improve its ability to synthesize Tyr derivatives using lower concentrations of phenol.

**Figure 2.**
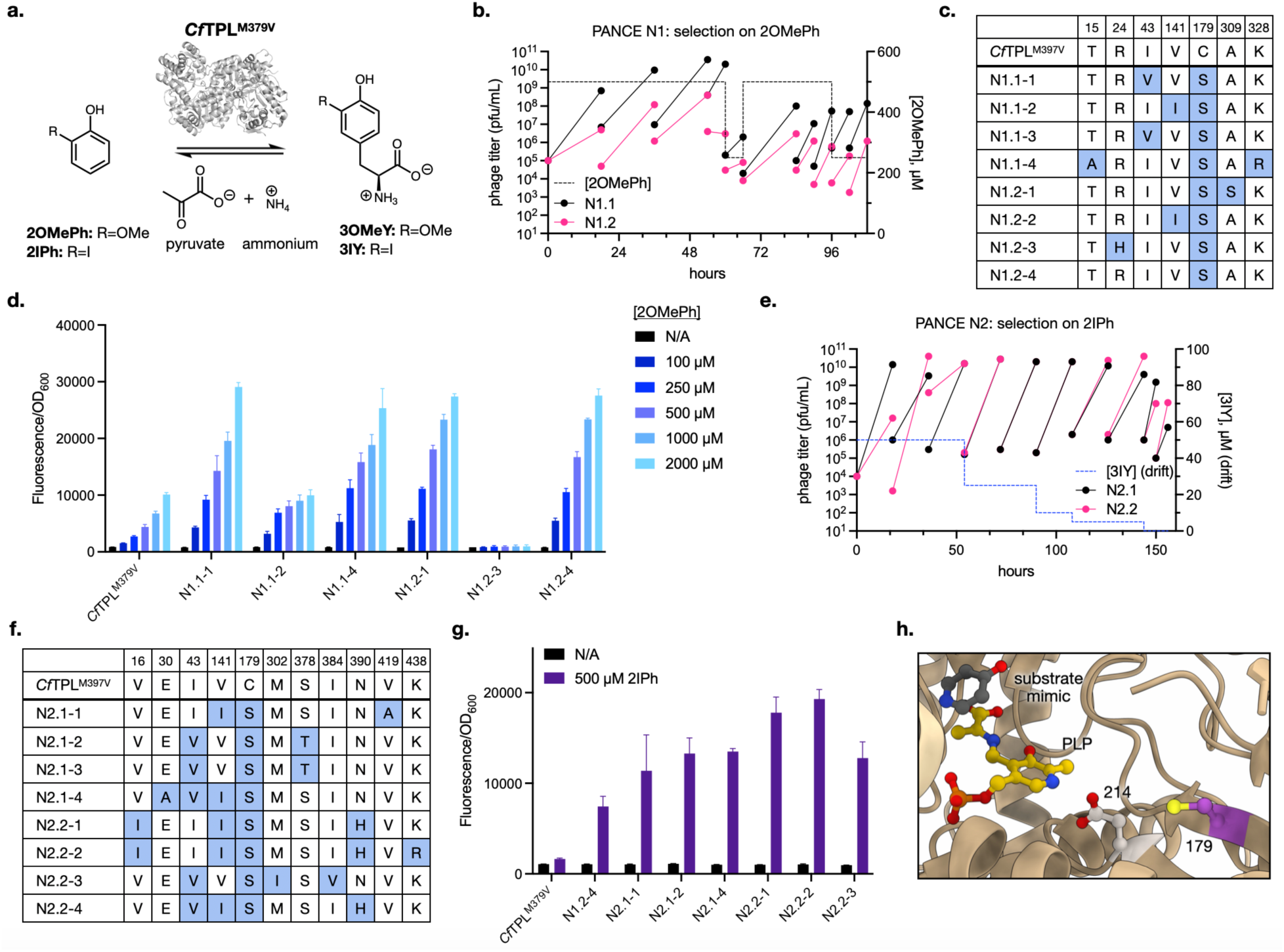
Tyrosine phenol lyase (TPL) evolution. **a.** *Citrobacter freundii* Tyrosine Phenol Lyase M379V (*Cf*TPL^M379V^) catalyzes the reversible condensation of 2-substituted phenols with ammonium and pyruvate to form 3-substituted tyrosines (PDB ID: 2TPL). **b.** Phage assisted noncontinuous evolution (PANCE) of *Cf*TPL^M379V^ for improved 3OMeY production using 2OMePh substrate (N1). Each passage is denoted by two points connected by a line, where the first point represents input phage titer and the second point indicates output titer. Two lineages (N1.1 and N1.2) were evolved in parallel using MP6 or MP4, respectively. **c.** Sequencing of four isolated clonal from each lineage of PANCE N1. **d.** Evaluation of *Cf*TPL^M379V^ variants from N1 for their ability to produce 3OMeY using varying concentrations of 2OMePh as a function of GFP* production. **e.** PANCE of the final N1 phage pools for 3IY production using 2IPh substrate. Two lineages (N2.1 and N2.2) were initiated using N1.1 and N1.2, respectively, and evolved in parallel. **f.** Sequencing of four isolated clonal from each lineage of PANCE N2. **g.** Evaluation of *Cf*TPL^M379V^ variants from N1 (N1.2-4) and N2 for their ability to produce 3IY using 0 (“N/A”) or 500 µM 2IPh according to GFP* production. **h.** Site of the C179S mutation mapped onto the crystal structure of the *Cf*TPL active site (PDB ID: 6MPD). Residue C179 is shown in purple, while D214 is in white, PLP in yellow, and substrate mimic in gray.

We first tested the ability of *Citrobacter freundii* Tyrosine Phenol Lyase M379V (*Cf*TPL^M379V^) to produce 3-methoxytyrosine (3OMeY) in *E. coli* from 2-methoxyphenol (2OMePh) by using an engineered *Mj*3OMeYRS/*Mj*tRNA^mut^ pair to incorporate the 3OMeY product into a sfGFP reporter with an amber stop codon (hereafter referred to as GFP*).^41^ *Cf*TPL^M379V^ is an engineered variant with broadened substrate recognition compared to wild-type *Cf*TPL.^36^ At 500 µM added 2OMePh, *Cf*TPL^M379V^ exhibited modest ability to produce 3OMeY as measured by GFP fluorescence (**SI Figure 2**). In contrast, GFP signal generated by cells supplemented with 500 µM racemic 3OMeY was > 3-fold higher.

We evolved *Cf*TPL^M379V^ using PANCE, a noncontinuous version of PACE that selects phage populations through serial passaging rather than continuous flow, to improve its ability to produce 3OMeY using reduced 2OMePh concentrations. Two lineages (N1.1 and N1.2) were evolved in parallel (**Figure 2b**). At the start of the experiment, phage were allowed to propagate overnight with 500 µM 2OMePh. Eventually, propagation time was decreased to 6 h and 2OMePh concentration lowered to 250 µM to increase selection stringency (**Figure 2b**). After 10 passages, we characterized four clonal phage from each lineage by sequencing. We observed universal convergence upon a C179S mutation, as well as multiple instances of I43V and V141I (**Figure 2c**). These clonal phage exhibited up to 1000-fold greater 2OMePh-dependent propagation activity compared to parent *Cf*TPL^M379V^ phage (**SI Figure 3**).

**Figure 3.**
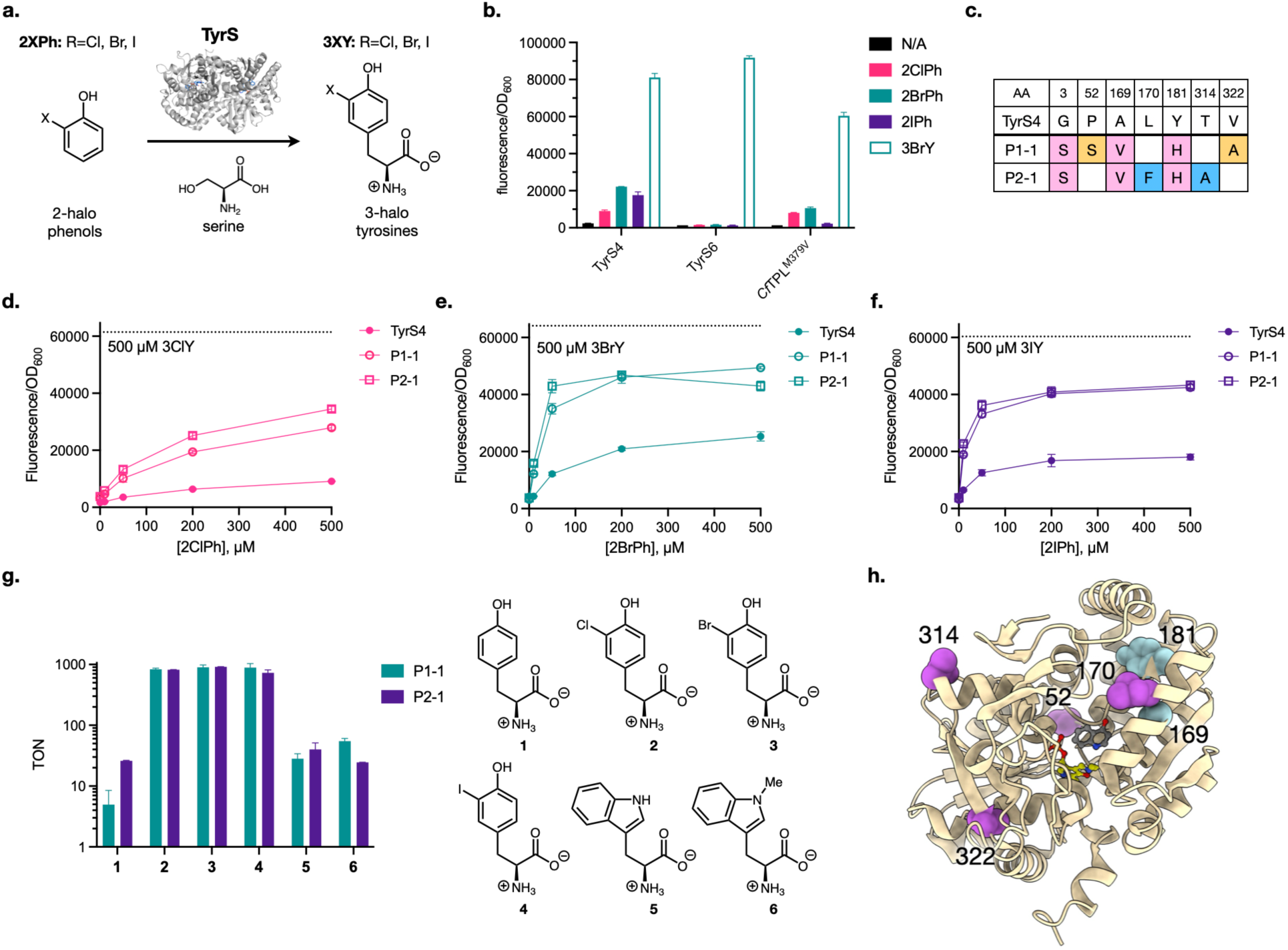
Phage assisted evolution improves the activity of TyrS4 on 2ClPh, 2BrPh, and 2IPh. **a.** TyrS enzymes catalyze the condensation of 2-halophenols (2XPhs) and serine to form 3-halotyrosines (3XYs) (PDB ID: 8EH1). **b.** Production of 3XYs from 2XPhs by TyrS4, TyrS6, and *Cf*TPL^M379V^ measured by incorporation into GFP*. **c.** Consensus mutations identified from PACE experiments using 2BrPh (P1-1) or 2IPh (P2-1)**. d.-f.** Evaluation of the production of **d.** 3ClY, **e.** 3BrY, or **f.** 3IY by evolved variants as a function of substrate concentration as measured by incorporation into GFP*. **g.** Analytical scale evaluation of the ability of P1-1 and P2-1 to synthesize amino acids **1**-**6**. The Turnover Number (TON) was determined using 0.1% of purified enzyme relative to the limiting substrate. **h.** Mutated residues highlighted in the crystal structure of TyrS1 in complex with PLP (yellow) and 4-hydroxyquinoline substrate mimic (gray) (PDB ID: 8EH1). Mutations in common between P1-1 and P2-1 are shown in blue; mutation unique to either variant are shown in purple. Position G3 is not resolved in the structure.

We subcloned the evolved *Cf*TPL^M379V^ variants onto plasmids and measured their ability to produce 3OMeY using varying concentrations of 2OMePh based on GFP* fluorescence (**Figure 2d**). Several of the evolved *Cf*TPL^M379V^ variants showed improvements, with the universally fixed C179S mutation alone conferring a > 2-fold increase in GFP signal for variant N1.2-4 compared to the parent enzyme. The N1.1-1 (I43V/C179S), N1.1-4 (T15A/C179S/K328R), and N1.2-1 (C179S/A309S) variants were also similarly improved in this experiment. In contrast, clone N1.1-2 (V141I/C179S) was no more active than the *Cf*TPL^M379V^ parent. However, cells expressing N1.1-2 also produced less GFP when supplemented with 3OMeY compared to cells expressing *Cf*TPL^M379V^ (**SI Figure 4**). Due to microscopic reversibility, any mutation that increases 3OMeY synthesis will also increase 3OMeY degradation, although the relative rates *in vivo* are determined by additional factors. We hypothesize that N1.1-2 is sufficiently activated that its rate of breakdown of 3OMeY product competes with the ability of 3OMeYRS to charge 3OMeY into GFP. Finally, R24H/C179S (N1.2-3) appeared to be inactive, and SDS-PAGE of cell lysates expressing N1.2-3 revealed that it has no detectable soluble expression, while the other evolved variants expressed at similar levels to the *Cf*TPL^M379V^ parent (**SI Figure 5**). Because clonal phage were sampled from a continuously mutagenized population, it is plausible that N1.2-3 had randomly acquired a deleterious mutation (R24H) prior to sampling. The incorporation of the biosynthesized 3OMeY into GFP was verified by mass-spectrometry (**SI Figure 6**).

**Figure 4.**
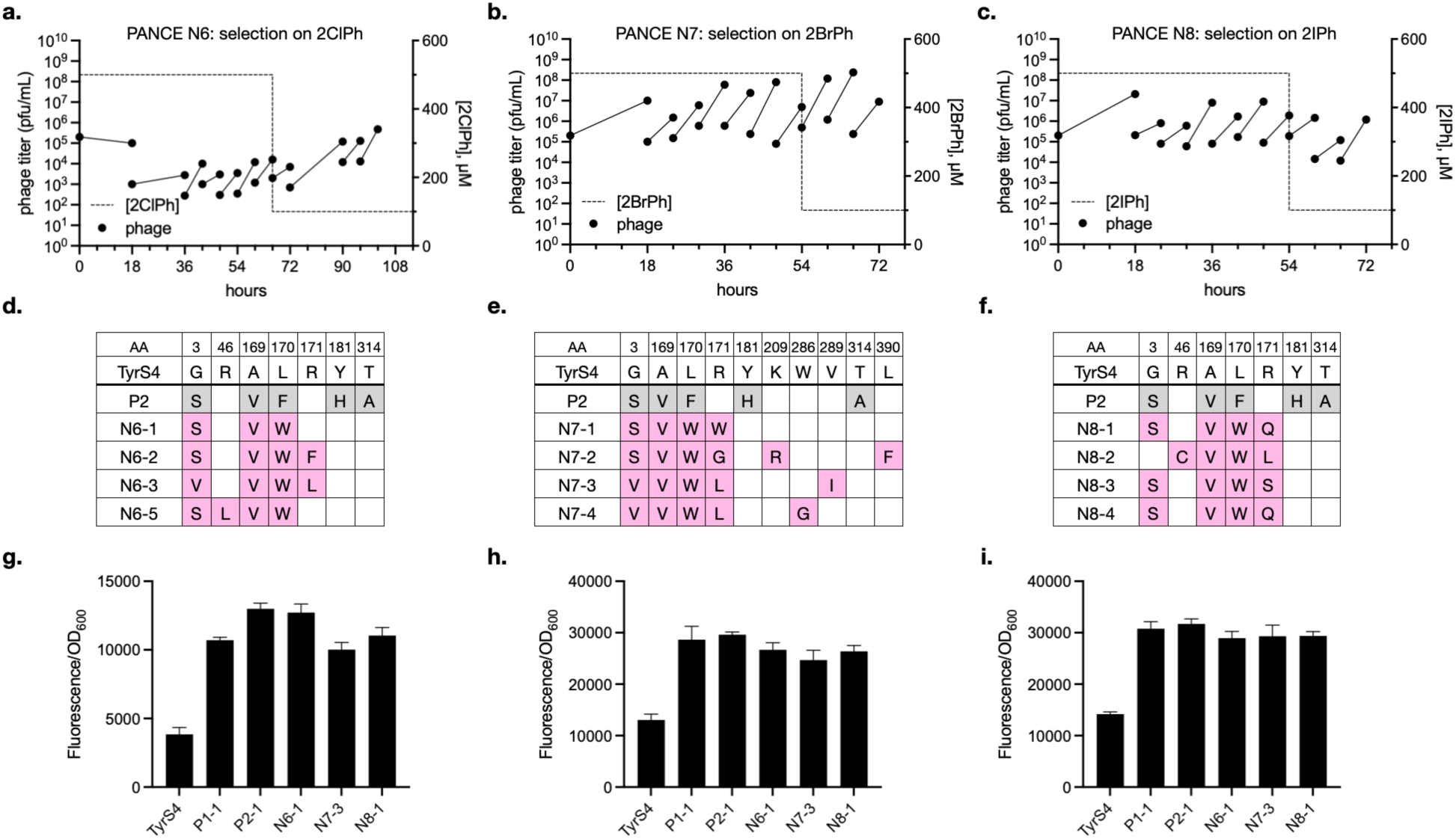
Selection of a TyrS4 site-saturation library. a.-c. A phage encoded library of TyrS4 variants with fully randomized residues at positions 169-171 was selected using *gIII\*\** AP using **a.** 2ClPh, **b.** 2BrPh, or **c.** 2IPh in the presence of additional mutagenesis by MP6. **d.-f.** Sequencing of four clonal phage from the last passage of selections shown in **a.-c.** Mutations shared with TyrS4 variant P2-1 are shown in pink and newly observed mutations shown in green. **g.-i.** Activity of plasmid-encoded TyrS4 variants in 3XY production from 50 µM **g.** 2ClPh, **h.** 2BrPh, and **i.** 2IPh measured by incorporation into GFP*.

**Figure 5.**
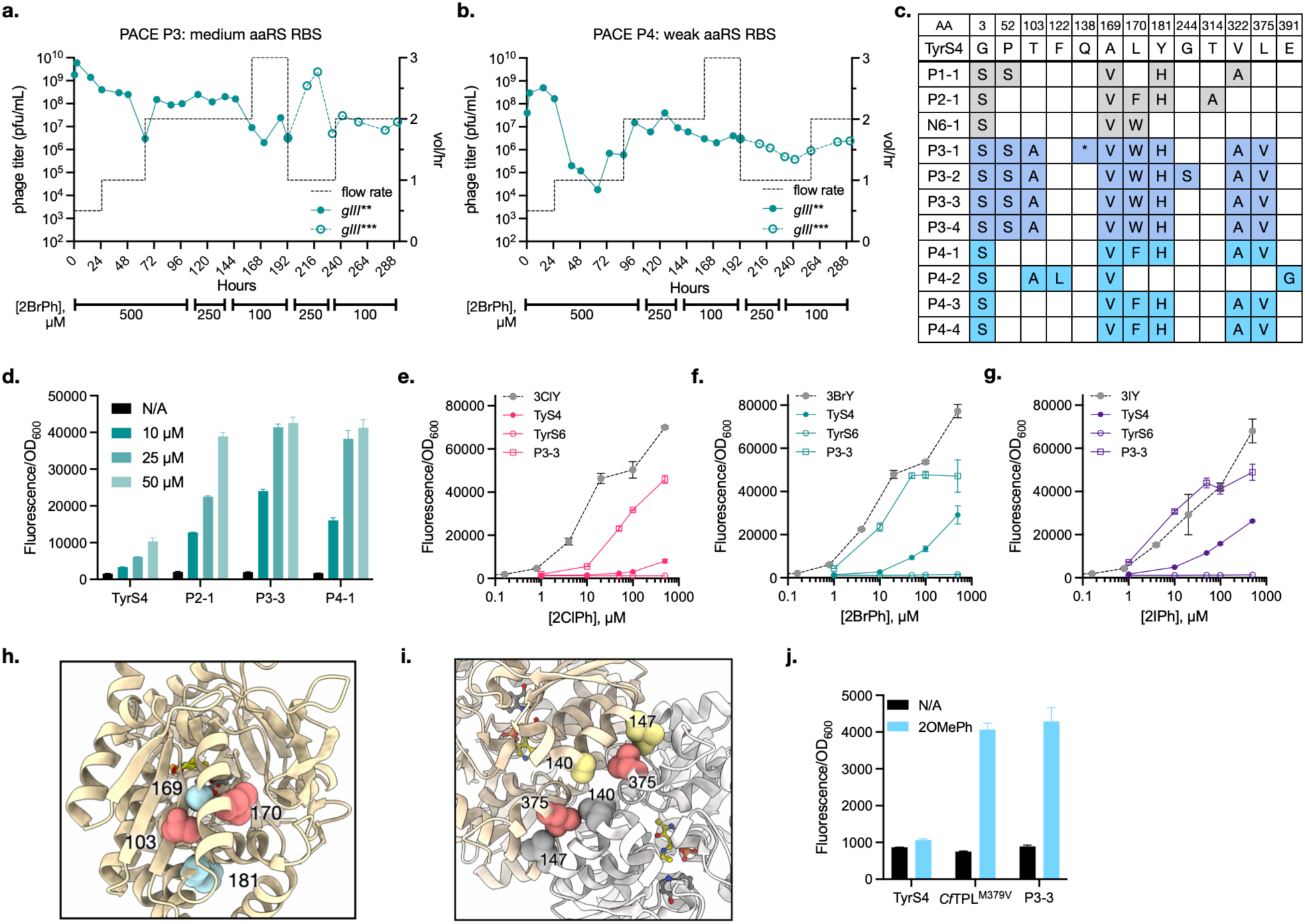
Reduced aaRS expression elicits further TyrS improvements in PACE. a.-b. PACE of TyrS using medium (**a.** P3) or weak (**b.** P4) ribosome binding sites for aaRS expression during selection. Selection initially utilized *gIII\*\**; after 192 hours, phage were evolved using *gIII\*\*\** to increase selection stringency. **c.** Sequencing of isolated phage from the final timepoint of PACE experiments P3 and P4. **d.** Comparison of 2BrPh conversion by TyrS variants TyrS4, P2-1, P3-3, and P4-1 measured by GFP* production. **e.-g.** GFP* fluorescence produced from titration of 2ClPh (**e.**), 2BrPh (**f.**), or 2IPh (**g.**) by TyrS4, TyrS6, or P3-3.

We further evolved the phage pool obtained from the 2OMePh PANCE to produce 3-iodotyrosine (3IY) from 2-iodophenol (2IPh), a substrate that TPLs have previously been shown to have no activity on.^39,43^ Here, we used an engineered *Mj*3XYRS/*Mj*tRNA^mut^ pair to incorporate 3IY into *gIII*\*\*.^44^ Because 2IPh is a more challenging substrate, we initiated PANCE with several drift passages by adding 3IY directly into the media, then gradually decreased the 3IY concentration through the course of the experiment. We performed the last two PANCE passages using shorter (6 h) propagation times with no added 3IY (**Figure 2e**). Sequencing of clonal phage isolated from the final passage showed several new mutations (**Figure 2f**). The clonal phage also displayed 100 to 1000-fold higher 2IPh dependent propagation relative to *Cf*TPL^M379V^ phage (**SI Figure 7**).

We tested the ability of plasmid-encoded *Cf*TPL^M379V^ variants to produce 3IY from 2IPh using the GFP* reporter (**Figure 2g**). All evolved variants showed notable 3IY generation with 500 µM 2IPh, as measured by GFP fluorescence, while *Cf*TPL^M379V^ was essentially inactive. The best variants, N2.2-1 and N2.2-2, produced > 10-fold greater fluorescence vs. *Cf*TPL^M379V^, ∼2-fold greater fluorescence vs. N1.2-4 (isolated from our initial PANCE using 2OMePh substrate), and generated ∼40% of the GFP signal observed from direct supplementation with 500 µM 3IY (**SI Figure 8**). The incorporation of the biosynthesized 3IY into GFP was verified by mass-spectrometry (**SI Figure 9**). SDS-PAGE analysis showed that the evolved variants had similar or modestly increased levels of expression compared to *Cf*TPL^M379V^ (**SI Figure 10**), suggesting that the gains observed on 2IPh are not primarily derived from higher enzyme expression. The 2IPh evolved variants also showed improved activity with 2OMePh, increasing 3OMeY production to a similar extent as the *Cf*TPL^M379V^ variants obtained from our first PANCE (**SI Figure 11**). However, they also showed moderately increased lyase activity compared to the parent enzyme, manifested as a reduction in GFP* fluorescence from direct 3OMeY supplementation (**SI Figure 4**).

The mutations enriched by our evolutions mainly lie outside of the *Cf*TPL active site. One exception is S378T, which is directly next to V379 (**SI Figure 12**). Prior mutation of methionine at position 379 to valine greatly increased the ability of *Cf*TPL^M379V^ to accept 2-substituted phenols; S378T may extend this effect to 2-iodophenol.^36^ The strongly enriched and highly activating mutation C179S is located ∼8 Å from the PLP cofactor and ∼5 Å from D214 (**Figure 2h**). The carboxylic acid moiety of D214 interacts with N1 of PLP and affects the electron withdrawing capacity of the cofactor.^45^ Replacement of C179 with serine, a stronger H-bond donor, could exert an influence on PLP through longer-range interactions. Finally, T15A, V16I, and I43V all localize to the interdimer interface of the *Cf*TPL tetramer (**SI Figure 13**). Mutations in this region in *Cf*TPL and other TPL homologs have been shown to increase protein stability.^46,47^ Notably, the T15A mutation was previously identified from screening a random mutagenesis library of *Cf*TPL variants for improved L-DOPA synthesis and was found to stabilize PLP binding.^46,47^ Taken together, these results demonstrate the ability of our selection strategy to identify improved variants of *Cf*TPL^M379V^ in less than two weeks.

### Continuous evolution of an engineered tyrosine synthase for improved halotyrosine synthesis

The tradeoff between improvements in ncAA biosynthesis and degradation we observed from our PANCE experiments with TPL led us to examine whether we could use our PACE selection to improve the ability of tyrosine synthases (TyrS), recently engineered from standalone tryptophan synthase *Tm*TrpB-9D8*,^32,48^ to biosynthesize tyrosine-derived ncAAs in *E. coli* from serine and phenols (**Figure 3a**). Similar to TPL, TyrS enzymes are PLP-dependent, but unlike TPL, TyrS does not exhibit lyase activity.^32^ We hypothesized that TyrS would represent a better starting point for engineering an enzyme to generate Tyr-derived ncAAs without requiring high concentrations of toxic phenols.

Although the engineered TyrS variants are reported to robustly synthesize tyrosine derivatives *in vitro*, efficient activity in a cellular context is a distinct selective outcome.^17,18^ Therefore, we tested plasmid-encoded TyrS variants TyrS4 and TyrS6 for their ability to biosynthesize 3-halotyrosines (3XYs) in *E. coli*.^32^ Cells expressing these enzymes were incubated with 2-halophenols (2XPhs) overnight using the engineered *Mj*3XYRS/*Mj*tRNA^mut^ pair to incorporate the 3XY product into GFP* (**Figure 3b**).^44^ For comparison, we also tested *Cf*TPL^M379V^. The most evolved TyrS variant, TyrS6, had little observable activity in this assay. A less-evolved enzyme, TyrS4, had a moderate increase in GFP fluorescence when 500 µM 2-bromophenol (2BrPh) or 2-iodophenol (2IPh) was supplemented, and a smaller increase when 500 µM 2-chlorophenol (2ClPh) was added. However, these GFP values were notably lower than those obtained when 500 µM 3BrY was added directly. Therefore, we elected to evolve TyrS4 to increase its ability to biosynthesize 3XYs in a cellular context.

Phage encoding TyrS4 showed poor fitness when tested in propagation assays using host cells carrying a AP with *gIII\*\** and the *Mj*3XYRS/*Mj*tRNA^mut^ pair (**SI Figure 14a**), consistent with the moderate activity we observed in the GFP* assay. Given their low starting propagation, we first evolved TyrS4 phage using PANCE in three separate lineages using 2ClPh, 2BrPh, or 2IPh as substrate (**SI Figure 14b-d**). Similar to our *Cf*TPL^M379V^ evolution, we initiated PANCE under less stringent conditions with overnight propagation and high substrate concentrations, then increased stringency by decreasing the propagation time or substrate, or both. After each lineage gained the ability to propagate under more stringent conditions, we transitioned them into PACE for further evolution (**SI Figure 15**). While phage selected using 2BrPh and 2IPh were eventually able to achieve high titers, phage selected on 2ClPh slowly declined over the course of the experiment, suggesting that they were unable to discover mutations enabling robust propagation under continuous flow conditions. Sequencing the 2BrPh and 2IPh populations after PACE revealed that the TyrS4 phage pools had strongly converged upon two similar but distinct genotypes (**Figure 3c**). Three mutations, G3S, A169V, and Y181H were shared between the 2BrPh and 2IPh phage pools, suggesting that these mutations are generally activating for 3XY synthesis by TyrS4. Additionally, Y181H and L170F, which fixed in the evolution using 2IPh, are also present in TyrS6.

When expressed from plasmids in *E. coli* and incubated with 2ClPh, 2BrPh or 2IPh, the variants isolated from evolution using 2BrPh (P1-1) and 2IPh (P2-1) showed markedly improved 3XY production, measured by GFP* suppression, compared to the TyrS4 parent (**Figure 3d-f**). Despite being evolved solely using a single substrate, P1 and P2 both exhibited a general increase in the ability to synthesize all three 3XYs, consistent with our hypothesis that the accrued mutations were generally activating rather than specifically enhancing activity on one substrate. 3XY biosynthesis through P1-1 and P2-1 enabled GFP* suppression to within ∼50-80% of values obtained through direct supplementation with the ncAA. Interestingly, GFP values began to saturate at ∼50 µM 2BrPh and 2IPh, suggesting that adding more substrate produces diminishing returns in GFP production. We confirmed using mass spectrometry analysis that GFP incorporating 3XY generated by P1-1 exhibited masses consistent with incorporation of each ncAA (**SI Figure 16**).

The evolved variants expressed to the same extent compared to TyrS4, suggesting that the observed activity improvements are not due to increases in solubility (**SI Figure 17**). We expressed and purified variant P1-1 and P2-1 (**SI Figure 18**) to test their activities *in vitro*. These enzymes showed robust ability to synthesize all three 3XYs, reaching the maximum number of turnovers over the duration of the experiment (**Figure 3g**). In contrast, they were less active in producing tyrosine or tryptophan from phenol or indole, respectively. Finally, we measured TyrS4, P1-1, and P2-1’s catalytic parameters for synthesizing 3BrY from 2BrPh (**Table 1** and **SI Figure 19**). Compared to TyrS4, P1-1 exhibited a ∼2-fold decrease in K_M_ and P2-1 showed improvements in both *k*_cat_ and K_M_.

**Table 1.** Properties of purified TyrS4 variants.

| Enzyme | $k_{cat}$ ($\text{min}^{-1}$) | $K_M$ 2BrPh ($\mu\text{M}$) | $k_{cat}/K_M$ ($\text{M}^{-1} \text{s}^{-1}$) |
| --- | --- | --- | --- |
| TyrS4 | $1.2 \pm 0.1$ | $336 \pm 23$ | $59.5 \pm 1$ |
| P1-1 | $1.2 \pm 0.1$ | $178 \pm 28$ | $112 \pm 5$ |
| P2-1 | $1.6 \pm 0.1$ | $187 \pm 26$ | $144 \pm 5$ |
| N6-1 | $1.4 \pm 0.1$ | $145 \pm 27$ | $161 \pm 7$ |
| N8-1 | $1.3 \pm 0.1$ | $133 \pm 27$ | $167 \pm 8$ |
| P3-3 | $0.4 \pm 0.1$ | $44 \pm 6$ | $152 \pm 40$ |

The mutations accrued in P1-1 and P2-1 map to several regions of the TyrS structure (**Figure 3h**).^32^ A169, L170, and Y181 reside in the COMM domain, whose rigid body open-close dynamics influence both the shape of the active site and the persistence of the reactive amino acrylate intermediate.^49,50^ Mutations throughout the COMM domain have often been observed in directed evolution of TrpB enzymes, including TyrS, but the identities of the beneficial mutations are highly variable and often specific to the selective pressure. Mutations G3, P52, V322, and T314 are distal to the active site, with G3 and T314 lying at solvent-exposed regions. Finally, three mutations, G3S, A169V, and Y181H are shared between both evolved variants.

To dissect the role of individual mutants in improving the activity of P1-1 and P2-1, we generated single reversions and tested their activities in the GFP assay (**SI Figure 20a-b**). These experiments revealed that the A169V mutation was the most important for the activity of both variants. We also examined the effects of individual mutations on the expression levels of P1-1 and P2-1 assessed through SDS-PAGE (**SI Figure 20c-d**). No single mutant appeared to exert a large effect except for A169V, whose impact was very different between the two variants. For P1-1, reversion of A169V caused a noticeable loss in expression, while in P2-1, this reversion appeared to boost protein production.

### Selection of a TyrS4 site-saturation library

The mutations enriched in the independent lineages of our evolved TyrS4 phage were highly similar, leading us to wonder if TyrS4 was limited to traversing a local fitness maximum by random MP-based mutagenesis. We therefore asked whether starting with a site-saturation library might enable more efficient exploration of the fitness landscape and potentially identify more active variants. We generated a phage-encoded library by performing triple saturation mutagenesis at A169/L170/R171 of the TyrS4 parent (theoretical size ∼35,000 members). These residues are located on a helix frequently mutated in directed evolution studies with TyrS and its TrpB parent;^23,51–54^ our own selections identified mutations at A169 and L170 that boosted TyrS4 activity.

We selected this library in parallel evolutions using 2ClPh, 2BrPh, or 2IPh substrate using PANCE. We began selection with low stringency (500 µM substrate), then decreased substrate to 100 µM after passage 7 to enrich more active sequences (**Figure 4a-c**). After a few passages, phage libraries selected with 2BrPh and 2IPh began to propagate robustly, while the library selected on 2ClPh exhibited lower fitness. Sequencing of four clonal phage from each final PANCE passage revealed that all three evolutions had once again converged on similar sequences (**Figure 4d-f**). At the region targeted by site-saturation mutagenesis, we observed universal enrichment of A169V and L170W, while mutations at position 171 were more divergent. In addition, we observed mutations at solvent-facing positions, including positions G3 and W286, which have been observed in previous evolution experiments with TrpB or TyrS4,^18,32^ that likely arose from random MP mutagenesis.

We tested the activities of three variants (N6-1, N7-3, and N8-1) identified from the library selection in the GFP* suppression assay (**Figure 4g-i**). Several of the new variants had comparable activity relative to P1-1 and P2-1 with all three 2XPhs. Given the fact that only 10-11 PANCE passages were required to identify these new variants (vs. > 13 passages in PANCE followed by > 150 hours of PACE), evolution using the multisite-saturation library was considerably more efficient. Assessment of a subset of variants by SDS-PAGE showed none of the tested mutants exhibited increased expression relative to the TyrS4 parent (**SI Figure 21**), suggesting that, similar to our PACE-evolved variants, the increases in activity are not due to solubility improvements. Additionally, we expressed and purified N6-1 and N8-1 and measured their catalytic parameters for 3BrY synthesis (**SI Figures 22 and 23**). Both variants exhibited slightly increased catalytic efficiencies compared to P1-1 and P2-1, driven by improvements in K_M_ (**Table 1**). These results show that PANCE of a modestly sized site-saturation TyrS4 library can greatly accelerate the identification of improved variants.

### Reduced aaRS expression enables evolution of highly active TyrS4 variants

Given the ability of our site-saturation library to identify improved TyrS4 variants with significantly less time invested in selection, we asked if the same strategy could improve the activity of PACE-evolved variant P2-1. We generated additional site saturation mutagenesis libraries based on P2-1 and selected them in PANCE at low (down to 10 µM) concentrations of 2BrPh or 2IPh. However, the resulting phage either did not converge on new genotypes or did not show improved activity (**SI Figure 24)**. In light of this, we surmised that that our current selection conditions could not enrich variants more active than P2-1 and reexamined our selection for ways to increase its stringency. We found that decreasing the strength of the aaRS ribosome binding site (RBS) decreased the propagation of phage carrying P2-1 with minimal impact to the selection’s operational range (**SI Figure 25**).

With more stringent selections in hand, we initiated PACE with the goal of improving upon P2-1. To maximize diversity of the starting population, post-PACE phage pools which yielded variants P1-1 and P2-1 as well as all previously selected site-saturation libraries were combined and used to infect two sets of lagoons distinguished by their *Mj*3XYRS expression levels (medium and weak RBS, respectively; **Figure 5a-b**). Lagoon flow rate, 2BrPh concentration, and the number of stop codons in *gIII* were modulated during the experiment to increase selective pressure. Sequencing of final phage populations showed strong convergence in each lagoon (**Figure 5c**).

Fluorescence produced from equivalent amounts of 3XY supplemented to cultures expressing an inactive TyrS is shown for comparison. **h.-i.** Mutated residues highlighted in the crystal structure of TyrS1 in complex with serine and 4-hydroxyquinoline (PDB ID: 8EH1). **i.** Residues mutated in P2-1 (A169 and Y181, light blue) and P3-3 (T103 and L170 orange). **j.** Mutated residue L375 (orange) sits at the TyrS dimer interface (protomers shown in wheat and white) and are near the TprB-9D8* mutation P140L and the TyrS4 mutation L147Q (positions shown as yellow and gray spheres). **j.** GFP* fluorescence produced by conversion of 2OMePh to 3OMeY by TyrS4, TPL^M379V^, or P3-3.

Phage evolved using moderate aaRS expression (P3) added mutations A103T, L170W, and L375V to the genotype of previously evolved variant P1-1. Phage evolved using weak aaRS expression (P4) also converged on L375V and A322V (which is also present in variant P1-1) in a genetic context that most closely resembles P2-1 with the notable omission of T314A. Variants P3-3 and P4-1 were examined for their ability to biosynthesize 3BrY from 2BrPh via incorporation into GFP* at low concentrations of 2BrPh (10-50 µM) (**Figure 5d**). Compared to P2-1, P3-3 and P4-1 produced similar levels of fluorescence from 50 µM substrate, but P3-3 and P4-1 achieved ∼2-fold increased fluorescence using 25 µM substrate. P3-3 also achieved ∼50% more fluorescence using 10 µM 2BrPh vs. P4-1, leading us to further characterize P3-3.

Titrating 2XPhs in the GFP* assay reveals P3-3’s activity at low substrate concentrations (**Figure 5d-f**), which has improved such that P3-3 achieves similar levels of fluorescence using 10 µM 2XPh compared TyrS4 supplemented with 50-fold more substrate. This observation is supported by our measurements of P3-3’s catalytic parameters (**SI Figures 26 and 27**), which revealed a dramatic (∼8-fold) reduction in K_M_ compared to TyrS4 parent, although at the expense of a ∼3-fold reduction in *k_cat_* (**Table 1**). Notably, at concentrations below 500 µM, the GFP fluorescence produced by P3-3 using 2BrPh or 2IPh is now comparable to directly supplementing cells with equivalent amounts of 3BrY and 3IY, respectively (**Figure 5f-g**).

Adding higher concentrations of 2BrPh or 2IPh, however, do not produce comparable levels of fluorescence as their tyrosine counterparts, suggesting that enzyme velocity relative to aaRS charging is still limiting under these concentrations. Additionally, although the fluorescence intensity from P3-3’s production of 3ClY from 2ClPh does not meet that of supplemented amino acid, P3-3 has the greatest improvement in activity on this substrate relative to the TyrS4 parent, producing ∼4-fold greater fluorescence at 500 µM 2ClPh, and > 9-fold at 50 µM 2ClPh (**Figure 5e**).

Several mutations gained in P3-3 are localized close to residues that were mutated to generate TyrS4 and TyrS6, as well as their predecessor *Tm*TrpB-9D8*. T103A and L170W, locations also mutated in TyrS4 and S6, respectively, join mutations we previously identified in or around the COMM domain, Y181H (also identified in TyrS6) and A169V, as well as preexisting TyrS4 mutations E105G, I128V N167D, I174T, and F184A (**Figure 5h**). These mutations may reshape the COMM domain to improve substrate binding. Moreover, mutations P52S and L375V exist at the TyrS homodimer interface, with L375V being directly opposite of 9D8* mutation P140L and TyrS4 mutation L147Q in the opposing protomer (**Figure 5i)**. Although the contributions of specific mutations are yet to be elucidated in future work, their locations are striking.

Finally, we examined the ability of P3-3 to generate 3OMeY using 2OMePh to see if increased activity on 2XPhs might extend to other 2-substituted phenols. While TyrS4 had no activity on 2OMePh, P3-3 produced comparable levels of GFP* fluorescence to *Cf*TPL^M379V^ (**Figure 5j**). Together, these results show that developing higher stringency selections enabled us to identify a more active TyrS variant using PACE.

## Discussion

Here, we demonstrate that phage-assisted continuous and noncontinuous evolution can be used to improve enzymes for *in situ* biosynthesis and incorporation of non-canonical tyrosines into proteins. Initially, we targeted *Cf*TPL^M379V^, a PLP-dependent C-C bonding forming enzyme that can produce substituted tyrosines from corresponding phenols. After twenty PANCE passages, we obtained *Cf*TPL^M379V^ variants that showed large increases in their production of 3OMeY from supplemented 2OMePh and furthermore acquired the ability to produce 3IY using 2IPh, a substrate that TPLs have no activity on.^39,43^ We then applied our selection to evolve a different enzyme: a tyrosine synthase, TyrS4, that was recently engineered from a standalone tryptophan synthase. We used a combination of PANCE and PACE to identify TyrS4 variants with improved ability to generate 3-halotyrosines using 2-halophenols.

Our most evolved variant, P3-3, possesses a ∼8-fold reduction in K_M_ for 2BrPh and exhibited substantial increases in 3XY production using lower phenol concentrations compared to TyrS4 parent. Finally, we show that P3-3 synthesized 3OMeY from 2OMePh with comparable activity to *Cf*TPL^M379V^, while TyrS4 was inactive on this substrate. Our results show that phage-assisted evolution can be used to improve existing enzyme function and to broaden enzyme activity to new substrates.

TPLs are widely employed for the synthesis of tyrosine-derived ncAAs in cells and *in vitro*. However, they require high concentrations of phenol substrate, which causes toxicity to cells and can inactivate the enzyme. Protein engineering can improve enzyme activity and stability, but there are only a handful of examples of engineered TPLs reported to date, with *Cf*TPL^M379V^ being arguably the most frequently used variant owing to its substrate tolerance. Our evolution of *Cf*TPL^M379V^ progressed rapidly, and we identified substantially activated variants within two weeks of evolution through PANCE. These experiments suggest that TPLs may be quite amenable to engineering and that PANCE can be used to quickly alter their substrate preferences and biosynthetic activities. Additionally, owing to their generally activated nature, the *Cf*TPL^M379V^ variants reported in this work may find utility in both biosynthetic and biocatalytic applications.

Recently reported tyrosine synthases were developed from the *Thermotoga maritima* tryptophan synthase β-subunit through *in vitro* directed evolution. Unlike TPL, TyrS enzymes do not possess lyase activity and may have increased utility for producing ncAAs in cells as they can in principle synthesize tyrosine derivatives using lower concentrations of phenols. However, we discovered that, of the reported tyrosine synthases, TyrS4 possessed only mild ability to biosynthesize 3XYs in cells, while the most advanced TyrS variant, TyrS6, was completely inactive, underscoring how enzymes optimized under *in vitro* conditions may require additional tuning prior to use in cells. Our evolution campaign of TyrS4 was more extensive than that of *Cf*TPL^M379V^, requiring both PANCE and PACE to identify activated variants P1-1 and P2-1 which were quite similar in genotype. We reconfigured our evolution strategy to include site saturation mutagenesis of TyrS4 in addition to MP-based random mutagenesis to see if improved variants could be identified more rapidly or with further improved activity. While this approach did yield activated variants in significantly less time, the variants exhibited comparable levels of activity to P1-1 and P2-1. Further improvements only emerged after we performed selections utilizing reduced aaRS expression, manifesting as greatly decreased K_M_ but at the cost of *k_cat_*. It is currently unclear to us if the trend of our evolutions primarily optimizing the K_M_ of TyrS variants can be attributed to the nature of our selection strategy or the evolvability of the target enzyme (or both). Developing strategies to specifically target *k_cat_* may be beneficial to future evolution campaigns.

Our study adds to a growing body of work employing phage-assisted evolution of enzymes for noncanonical amino acid biosynthesis.^30,31^ We extend this method to include three more engineered aaRS/tRNA pairs and demonstrate the evolution of two enzymes for improved 3-substituted tyrosine biosynthesis via C-C bond forming reactions. These efforts have broadened the range of ncAAs that can be cheaply biosynthesized and have demonstrated the modularity of phage-assisted evolution selections towards diverse target ncAAs. We expect our work to enable further implementation of genetic code expansion in biotechnological applications as well as future efforts to engineer enzymes for ncAA production in cells.

## Materials and Methods

### General methods

Antibiotics (Gold Biotechnology) were used at the following working concentrations: ampicillin, 50 µg/mL; spectinomycin, 100 µg/mL; kanamycin, 50 µg/mL; chloramphenicol, 25 µg/mL. Nuclease free water (Omega Biotek) was used for PCR reactions and cloning. For all other experiments, water was purified using a MilliQ purification system. Q5U DNA polymerase (New England Biolabs) or Q5 polymerase (New England Biolabs) was used for PCRs. A full list of plasmids used in this work is provided in **Supplementary Table 1**. Key primers are given in **Supplementary Table 2**. A full list of reagents and equipment used in this work is given in **Supplementary Table 3**. DNA sequences used in this work are given in **Supplementary Table 4**. Plasmid maps in GenBank format are provided in **Supplementary Data 1**.

Strain S2060 was used for plasmid cloning, amplification, and fluorescence assays. Strain S2208 was used for SP cloning and plaque assays.^55^ S2060 Δ*tnaA* Δ*trpAB* was generated by the lambda red recombineering method as previously reported^56^ and was used for all phage propagation assays and evolution experiments. Unless otherwise noted, cultures were grown at 37 °C shaking at 250 rpm. Plasmids and SPs were cloned by USER assembly or blunt-end ligation. Plasmids and SPs were sequenced using the SAVEMONEY strategy.^57^ Competent cells were prepared and transformed using the TSS method.^58^ Phage propagation and plaque assays were performed as previously described.^26^ Unless otherwise noted, phage propagation assays used an input of 10^5^ phage to infect a 2 mL culture of host cells for 6 h prior to output phage collection.

### PANCE

PANCE experiments were performed as previously described.^26^ S2060 Δ*tnaA* Δ*trpAB* containing the desired accessory plasmid, genetic code expansion plasmid, and mutagenesis plasmid (MP4 or MP6) were grown overnight in Davis Rich Media (DRM) with maintenance antibiotics.^59^ These cells were diluted 1:50 into DRM with maintenance antibiotics, grown to exponential phase, infected and supplemented with 500 µM of the desired phenol (unless otherwise noted) and 1 mM arabinose to induce the mutagenesis plasmid, and propagated for six hours or overnight. Cultures were then pelleted and the supernatant, containing phage, stored at 4 °C.

### PACE

PACE experiments were performed as previously described.^26^ Several colonies of freshly transformed S2060 Δ*tnaA* Δ*trpAB* containing the appropriate accessory plasmid, genetic code expansion plasmid, and MP6, were added to 1 mL of DRM, vortexed, and serially diluted 1:5 into DRM with maintenance antibiotics. All serial dilutions were grown overnight in a 96-well plate covered with porous film. The following day, serial dilutions at exponential growth phase were diluted into 8 mL DRM with maintenance antibiotics and grown to exponential phase. This culture was used to inoculate a 40 mL chemostat and was grown to exponential phase before beginning continuous flow. Chemostat flow rates began at 30 mL/hour and were adjusted to maintain cells at exponential growth (approximate OD_600_ values of 0.2 - 0.8). A syringe pump (New Era Pump Systems Inc.) supplied lagoons with solution containing arabinose (to induce MP6 mutagenesis), substrate, and ethanol (to solubilize substrate). Syringes were prepared at 50x initial intended working concentrations, with the flow rate being subsequently adjusted to reduce working substrate concentrations. For an initial lagoon working concentration of 2 mM arabinose and 500 µM substrate, syringes were loaded with 50 mL of solution containing 100 mM arabinose, 25 mM phenol, and 5% ethanol. Before phage infection, 15 mL lagoons were filled from the chemostats and given a bolus of 300 µL inducer/substrate solution and then equilibrated for one hour with a lagoon flow rate of 15 mL/hr and a syringe flow rate of 300 µL/hr. At the time of phage infection, lagoon flow rate was reduced to 7.5 mL/hr and syringe flow rate to 150 µL/hr. Lagoon and syringe flow rates were subsequently adjusted as desired over the course of the experiment. Lagoons were sampled twice daily by drawing 1 mL through the waste needle of each lagoon.

### GFP* plate assays

S2060 cells transformed with amber-suppressed sfGFP reporter (GFP*) plasmid, genetic code expansion plasmid, and ncAA synthase plasmid were inoculated into 1 mL 2xYT with maintenance antibiotics in triplicate and grown overnight. The following day, cultures were diluted 1:100 into DRM with antibiotics in 96-well plates. Cultures were grown to exponential phase (approximately three hours) before adding inducer and substrate. 100x inducer/substrate solutions were prepared to achieve final concentrations of 1 mM arabinose (to induce ncAA synthase), 1 mM IPTG (to induce GFP*), 100 ng/mL anhydrotetracycline (aTc, to induce genetic code expansion machinery), and the substituted phenol or tyrosine at the desired concentration. Inducer/substrate solutions were supplemented with 40% ethanol to ensure phenol solubility, resulting in 0.4% ethanol in bacterial culture. After induction, cultures were grown overnight for 19-21 hours. 100 µL of each culture was transferred to a black, clear-bottom 96 well plate and fluorescence and cell density measured in a Tecan Infinite M plex plate reader. Fluorescence values were measured with gain set to 80.

### SDS-PAGE of enzyme variants

300 μL of each triplicate culture from GFP* plate assays described above were combined, normalized by OD_600_, and pelleted by centrifugation for 2 min at 8,000 rcf. Supernatant was removed and pellets were resuspended in 150 μL BPER-II and incubated for 10 min at room temperature. Lysates were then pelleted for 10 min at maximum centrifuge speed at 4 °C. 10 µL of the soluble fraction was combined with 20 µL nuclease free water and 10 µL 4x Laemmli sample buffer (BioRad) containing 100 mM DTT or 1.4 M ϕ3-mercaptoethanol. Samples were incubated for 10 minutes at 95 °C and then loaded onto a 4-15% polyacrylamide gel (BioRad). Proteins were separated at 180-220 V.

### GFP purification

2 mL of 2xYT with maintenance antibiotics was inoculated with S2060 cells containing plasmids encoding GFP*, aaRS/tRNA pairs, and synthase variants and grown overnight. Overnight cultures were diluted 1:100 into 15 mL DRM containing maintenance antibiotics and grown for three hours. Cultures were then induced with 100-200 ng/mL aTc ,1 mM arabinose, and 1 mM IPTG and supplemented with either phenol substrate (200-500 µM) or tyrosine-derived ncAA (500 µM). After 14 hours of further growth, cultures were pelleted (20 min, 4,000 rcf, 4 °C) and resuspended in 1 mL room temperature B-PER II and lysed for one hour on ice, inverting every ten minutes. Lysates were pelleted for 20 min at 20,000 rcf and 4 °C. Lysates from duplicate cultures were combined, and each supernatant was mixed with 100 µL Ni-Sepharose resin (Cytiva) that was washed thrice in TE7 buffer (10 mM Tris, 1 mM EDTA, pH 7.5) and incubated for 90 minutes at 4 °C with head-to-tail inversion. The resin was pelleted (5 min, 900 rcf, 4 °C) and washed three times with 1 mL ice cold TE7 buffer containing 25 mM imidazole. To elute bound proteins, resin was resuspended in 100 µL cold TE7 with 250 mM imidazole, incubated for 2 minutes on ice, and pelleted. Imidazole was removed from the eluted protein using Zeba spin columns (Thermo).

### Mass spectrometry analysis

Agilent 6230B time of flight (TOF) mass spectrometer was used for electrospray ionization liquid chromatography mass spectrometry (ESI-LC-MS). Intact protein samples of GFP incorporated with ncAAs (5 µM, 2 µL injection) were separated on a PLRP-S 1000 Å, 50 ×1 mm, 5 μm column at 80 °C using a 3.9-minute gradient from 20 to 60% B (A=0.1% formic acid in water, B=acetonitrile). The maximum entropy charge deconvolution algorithm in Agilent MassHunter BioConfirm v10.0 was used to determine the neutral mass of intact proteins.

### Protein expression and purification

Individual colonies of BL21(DE3) cells transformed with synthase plasmid were inoculated into 2 mL 2xYT with kanamycin and grown overnight. The cultures were diluted 1:100 in 2xYT with maintenance antibiotics and were grown at 37 °C until they reached an OD_600_ of 0.7. 1 mM of Isopropyl *β*-D-1-thiogalactopyranoside (IPTG) was used to induce the cultures which were then incubated at 30 °C for a further 18 hours. Cultures were harvested by centrifugation (20 min, 4300 rcf), and the cell pellets were stored at -80 °C until purification.

Pellets were thawed on ice followed by resuspension in 7mL of Lysis Buffer (25 mM potassium phosphate pH 8.0, 100 mM NaCl, 20 mM imidazole, 100 µM PLP, 1 mM phenylmethylsulfonyl fluoride (PMSF), 2 mM Benzamidine, 0.02 mg/mL DNAse I, and 1 mg/mL lysozyme) and sonication (1” pulse, 1” pause, 30% Amp for 1 minute and repeated 4 times). The lysate was then incubated in a water bath at 75 °C (for TyrS4) or 68 °C (for all evolved variants) for 10 minutes to precipitate *E. coli* proteins. The heat-treated lysate was then clarified by centrifugation (30,000 rcf for 30 minutes at 4 °C) and the supernatant (Sup) was incubated with 1 mL of Ni-Sepharose resin (Cytiva) for 2 hours at 4 °C with gentle agitation. The resin was first separated from the flow through (FT) via centrifugation and then washed with 10 mL of wash buffer (25 mM potassium phosphate pH 8.0, 100 mM NaCl, and 50 mM imidazole) and bound proteins were eluted with 5 mL of elution buffer (25 mM potassium phosphate pH 8.0, 100 mM NaCl, and 300 mM imidazole). The eluted fraction was incubated with 1 mM of PLP for 18 hours at 4 °C, then buffer exchanged into 50 mM potassium phosphate buffer (pH 8.0) and concentrated to the desired protein concentration using Amicon Ultra-15 Centrifugal Filters (Millipore). The protein concentration was determined by Braford assay (Thermo).

### Analytical scale reactions and enzyme kinetics

Small-scale (100 µL) reactions to measure the TONs of the enzymes for different substrates were performed with 10 µM enzymes and 10 mM substrates in a buffered reaction mixture containing 10 mM L-Serine and 1 mM PLP in 50 mM potassium phosphate buffer (pH 8.0). The reactions were performed at 37 °C and were quenched after 18 hours, with 400 µL of the quenching solution containing 50% acetonitrile and 0.5 M HCl. The products were analyzed by reverse phase chromatography using either an Intrada Amino Acid column (Imtakt) or a BEH C18 column (Waters). 3 µL injections were made on a Acquity UHPLC (Waters) with an Acquity QDA MS detector (Waters).^13^

For Michaelis-Menten analysis, the 200 µL reactions were performed in similar manner and were quenched after 3 hours with 200 µL of quenching solution containing 80% acetonitrile and 0.8 M HCl. The products were analyzed by reverse phase chromatography with a BEH C18 column (Waters). 10 µL injections were made on a Acquity UHPLC (Waters) with an Acquity QDA MS detector (Waters).

## Author Contributions

J.S.A. and A.B. designed research, conducted experiments, and analyzed data. D.D. and A. M. W. designed and performed intact protein mass spectrometry experiments, data analysis, and interpretation. A.R.B. supervised research. T.W. designed and supervised research. All authors wrote the manuscript.

## Acknowledgements

The authors are grateful to Reece Gardner for guidance with LCMS experiments. This work was supported by the Office of the Vice Chancellor for Research and Graduate Education at the University of Wisconsin-Madison with funding from the Wisconsin Alumni Research foundation. J.S.A and A.B. were supported by NIH grants R35 GM159851 (to T.W.) and R35 GM153276 (to A.R.B). J.S.A was also supported by the Steenbock Predoctoral Graduate Fellowship administered by the University of Wisconsin-Madison Department of Biochemistry. A.R.B. additionally acknowledges the support of the Alfred P. Sloan Foundation. A.M.W. acknowledges the support of a David and Lucille Packard Fellowship for Science and Engineering (2021-73007) and a Career Award at the Scientific Interface from the Burroughs Wellcome Fund (1017065).

