## Supplementary Information for "Phage-assisted continuous evolution of enzymes for noncanonical tyrosine biosynthesis"

**Table of contents:**

**Supplementary Figures 1-27**

**Supplementary References**

**Supplementary Figures**

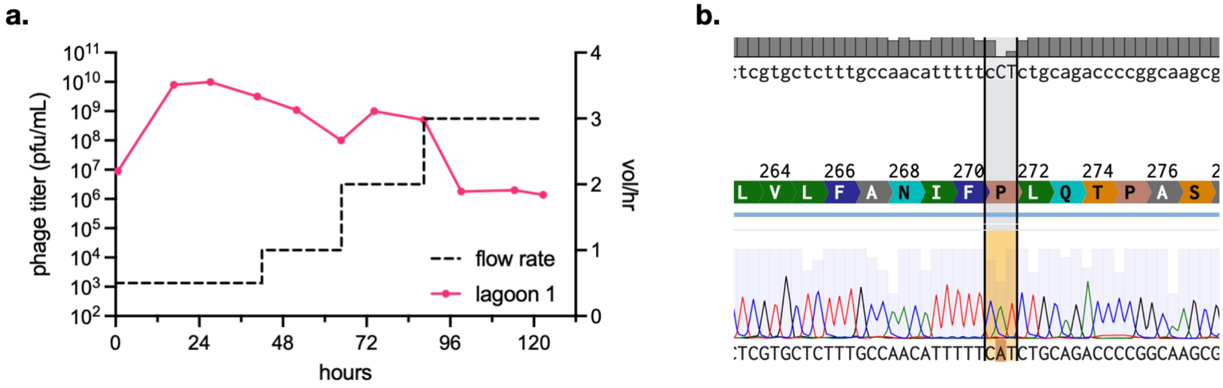


**Supplementary Figure 1. a.** PACE of phage encoding MfnG H271P using host cells carrying an accessory plasmid encoding *gIII***, *Mj*OMeYRS/*Mj*tRNA$^{mut}$, and MP6. **b.** Sanger sequencing of the phage pool after 24 hours of PACE show the majority of the population reverts the H271P mutation.

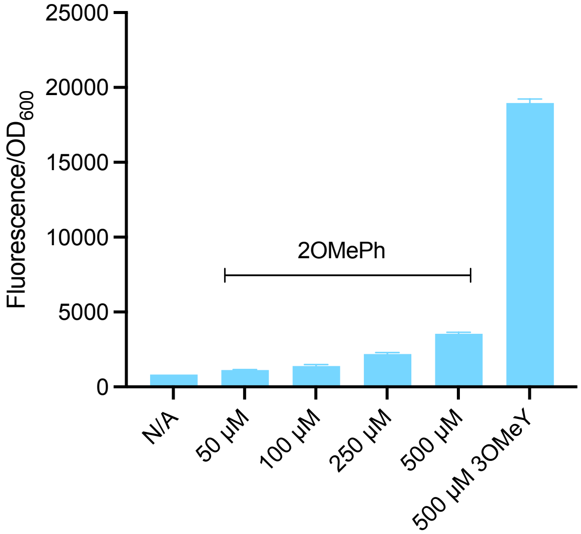


**Supplementary Figure 2.** *Cf*TPL$^{M379V}$ displays modest activity towards 2OMePh. Cultures expressing *Cf*TPL$^{M379V}$, *Mj*3MeOYRS/tRNA, and GFP* were cultured overnight with varying concentrations of 2OMePh or 500 $\mu$M racemic 3OMeY.

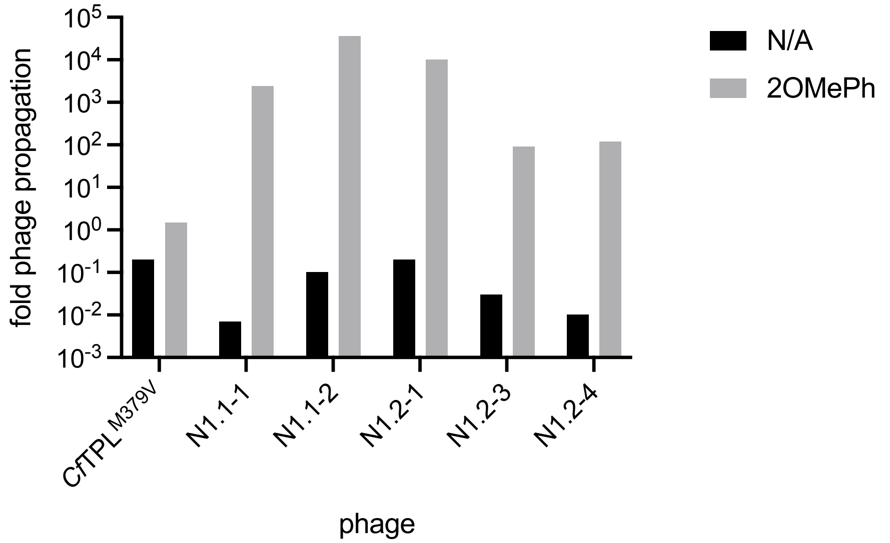


**Supplementary Figure 3.** Propagation of clonal phage isolated from PANCE of *Cf*TPL$^{M379V}$. Phage were propagated with either no substrate ("N/A") or 500 $\mu$M 2OMePh.

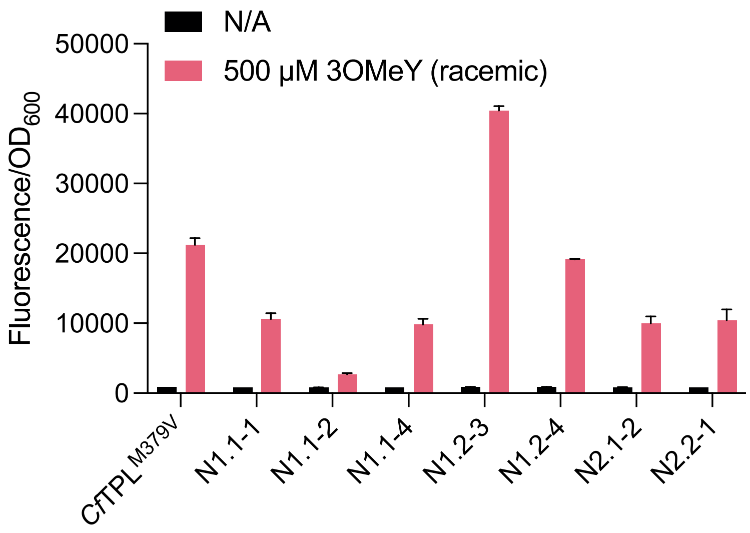


**Supplementary Figure 4.** Evaluation of *Cf*TPL$^{M379V}$ variants for 3OMeY decomposition as measured by 3OMeY incorporation into GFP*. Cultures expressing parent, or N1 or N2 *Cf*TPL$^{M379V}$ variants, *Mj*3OMeYRS/tRNA, and GFP* were cultured overnight with 0 $\mu$M ("N/A") or 500 $\mu$M racemic 3OMePh. Increased 3OMeY decomposition manifests as reduced GFP production and fluorescence.

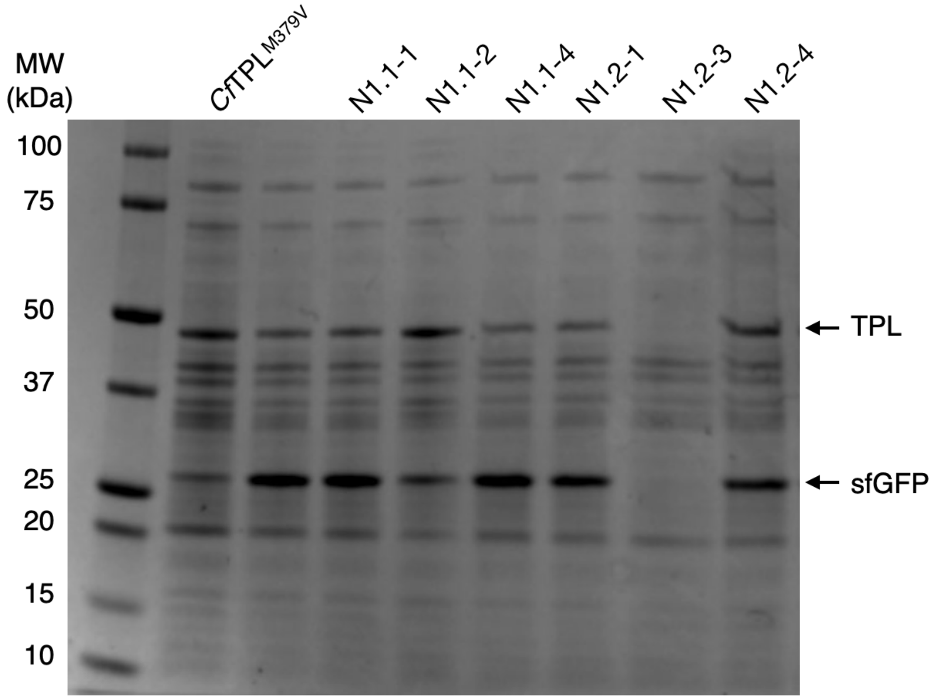


**Supplementary Figure 5.** Expression of *Cf*TPL$^{M379V}$ variants evolved from PANCE N1.

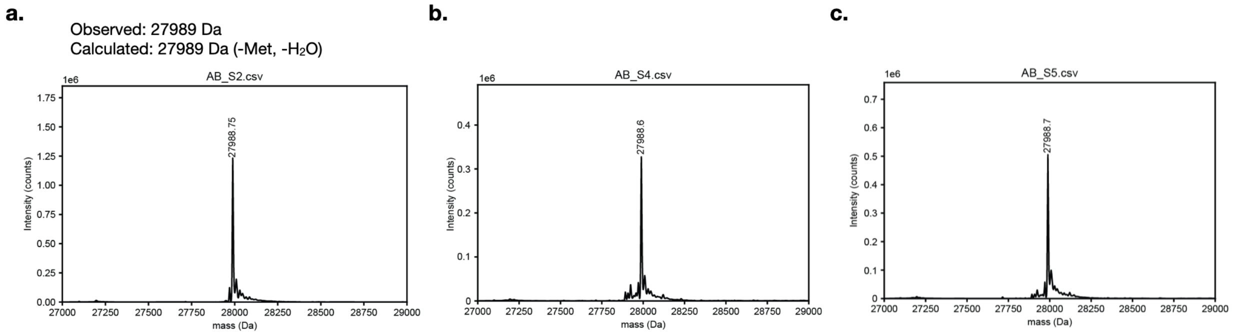


**Supplementary Figure 6.** Mass spectrometry analysis of GFP* produced in the presence of 500 $\mu$M **a.** 3OMeY, **b.** 2OMePh and N1.2-4 or **c.** 2OMePh and N2.2-1. Observed masses are consistent with incorporation of 3OMeY along with removal of the *N*-terminal Met and the loss of a water molecule from fluorophore maturation.

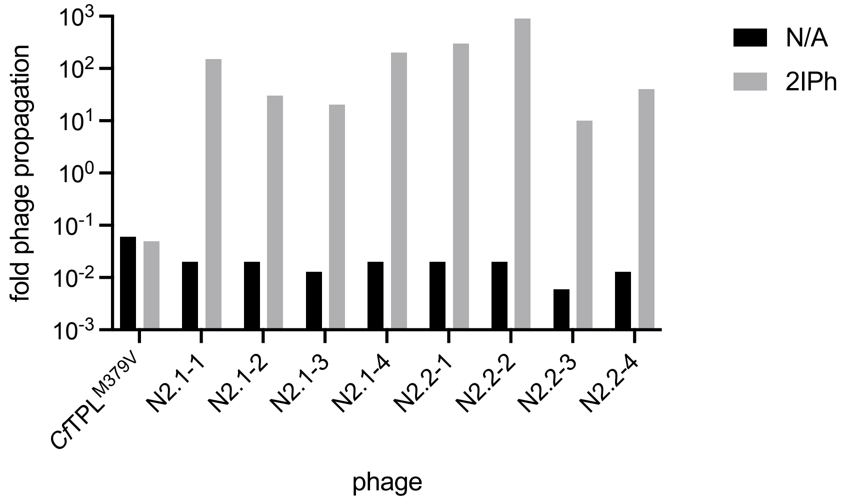


**Supplementary Figure 7.** Propagation of clonal phage isolated from PANCE of *Cf*TPL$^{M379V}$. Phage were propagated with either no substrate ("N/A") or 500 $\mu$M 2IPh.

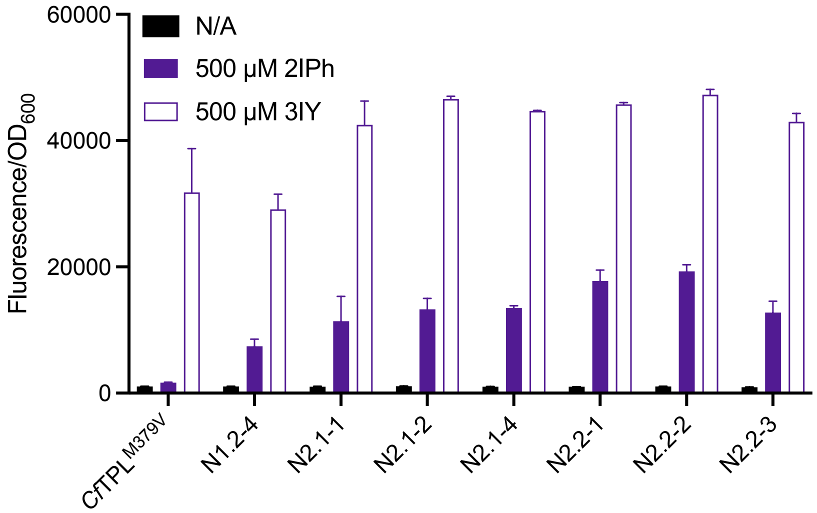


**Supplementary Figure 8.** Evaluation of PANCE N2 *Cf*TPL$^{M379V}$ variants for 3IY biosynthesis and incorporation into GFP*. Cells expressing *Cf*TPL$^{M379V}$ variants, *Mj*3XYRS/tRNA, and GFP* were cultured overnight with 0 $\mu$M ("N/A") or 500 $\mu$M 2IPh, or 500 $\mu$M 3IY.

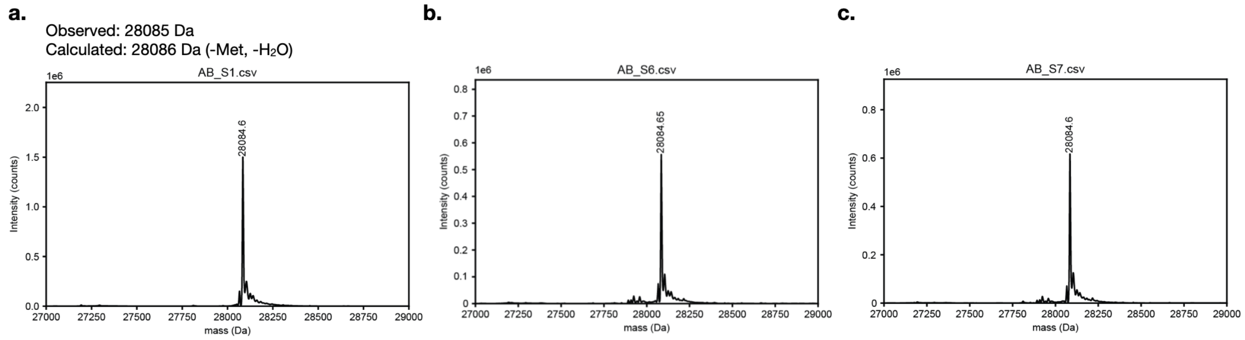


**Supplementary Figure 9.** Mass spectrometry analysis of GFP* produced in the presence of 500 $\mu$M **a.** 3IY, **b.** 2IPh and N1.2-4 or **c.** 2IPh and N2.2-1. Observed masses are consistent with incorporation of 3IY along with removal of the *N*-terminal Met and the loss of a water molecule from fluorophore maturation.

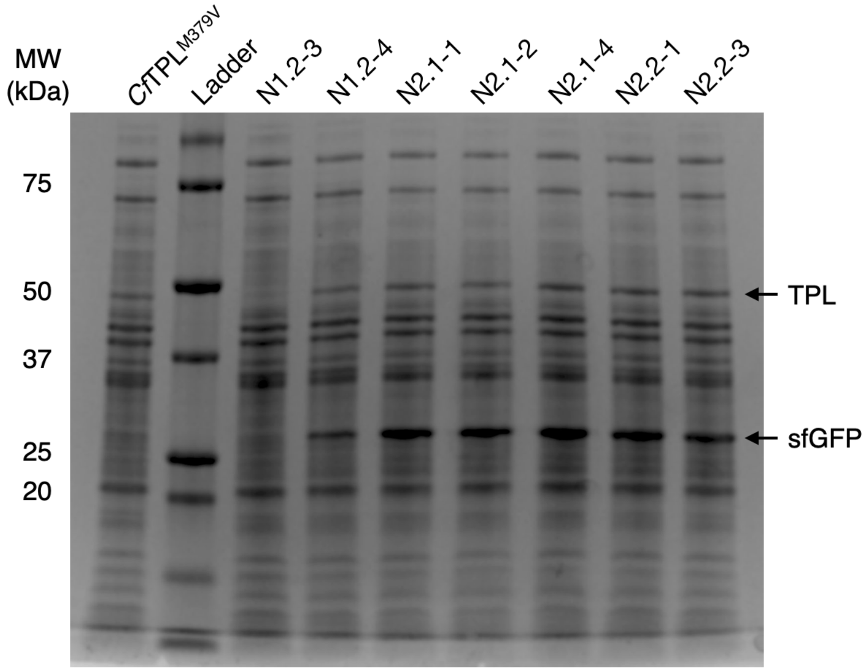


**Supplementary Figure 10.** Expression of variants from PANCE N1 and N2 in comparison to parent *Cf*TPL$^{M379V}$, assessed by SDS-PAGE.

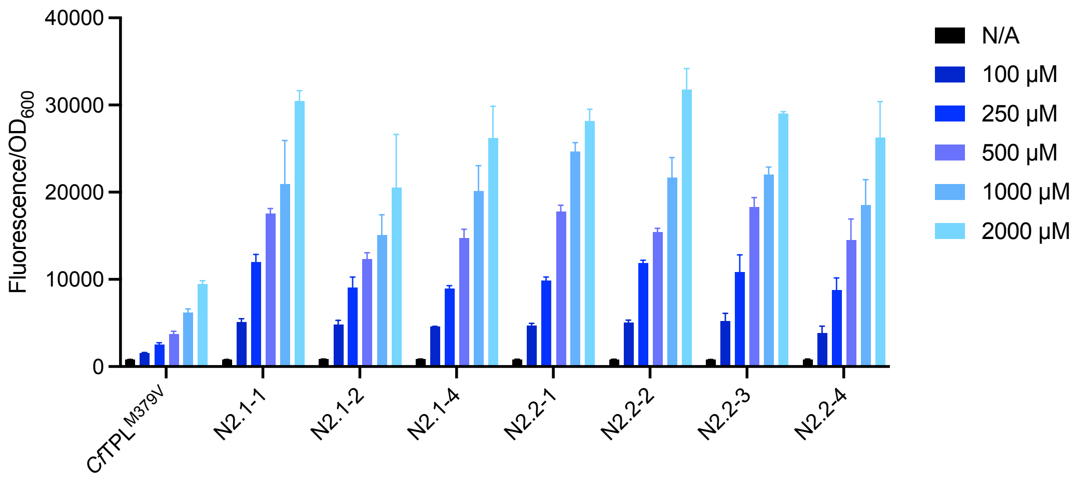


**Supplementary Figure 11.** Evaluation of PANCE N2 *Cf*TPL$^{M379V}$ variants for 3OMeY biosynthesis and incorporation into GFP*. Cells expressing *Cf*TPL$^{M379V}$ variants, *Mj*3OMeYRS/tRNA, and GFP* were cultured overnight with 0 *μ*M (“N/A”) or varying concentrations of 2OMePh.

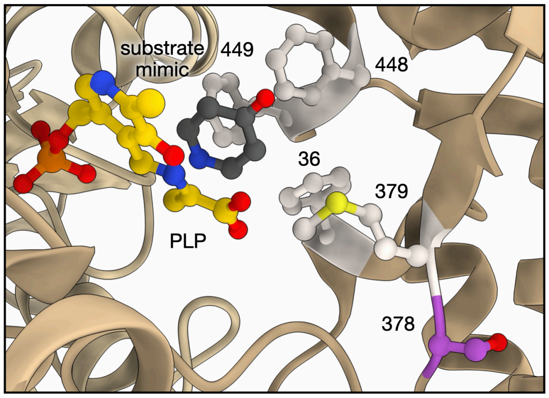


**Supplementary Figure 12.** Location of S378 (purple) mutation with respect to the active site of *Cf*TPL (PDB ID: 6MPD). PLP is shown in yellow and substrate mimic in gray; residues F36, M379, F448 and F449 are shown in white.

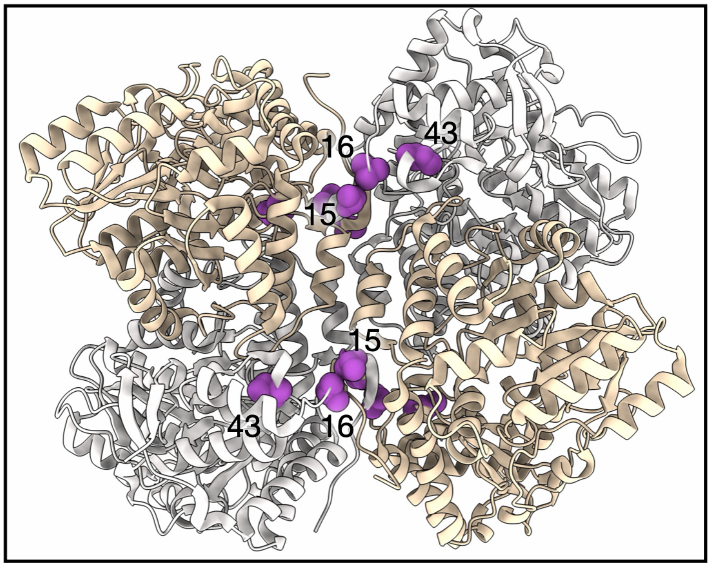


**Supplementary Figure 13.** Location of T15, V16, and I43 mutations (purple) at the interface the TPL homo-tetramer (PDB ID: 7TDL).

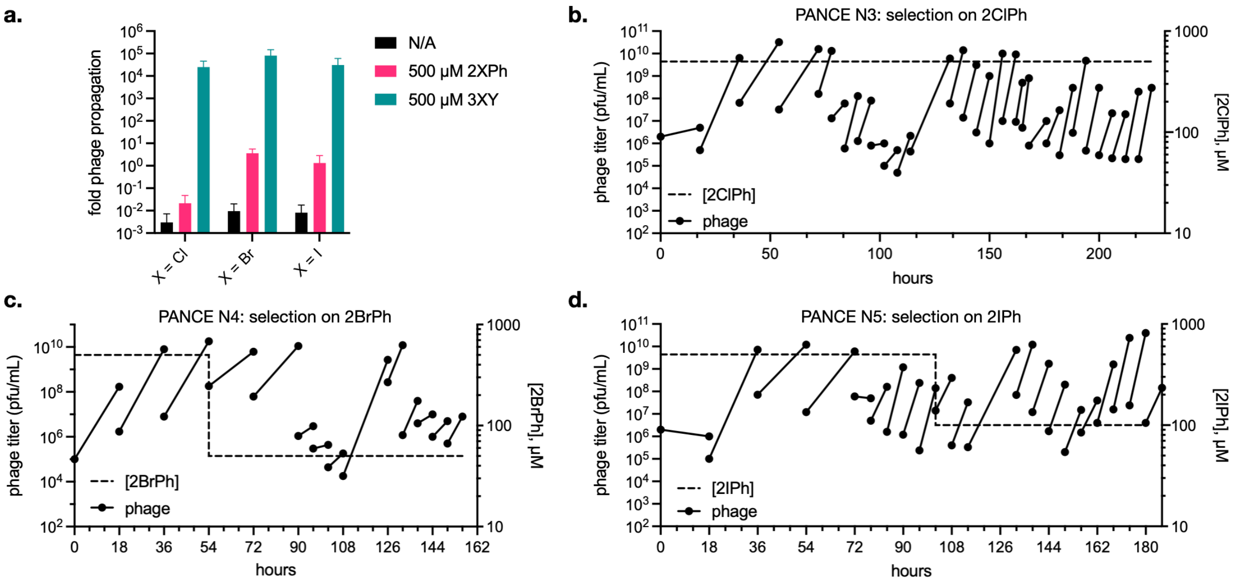


**Supplementary Figure 14.** PANCE of TyrS4 for the formation of 3ClY, 3BrY, or 3IY. **a.** 6-hour propagation of phage encoding TyrS4 on host cells with the *gIII*** accessory plasmid with supplementation with 500 $\mu$M 2XPh, 3XY, or no substrate ("N/A"). **b.-d.** Evolution of TyrS4 phage in PANCE with 500 $\mu$M **b.** 2ClPh, **c.** 2BrPh, or **d.** 2IPh. 2XPh concentrations were reduced in later passages to 100 $\mu$M (2BrPh and 2IPh) or 50 $\mu$M (2BrPh) to further increase selective pressure. Each passage is represented by two points connected by a line, where the first point signifies the input phage titer, and the second point the output phage titer.

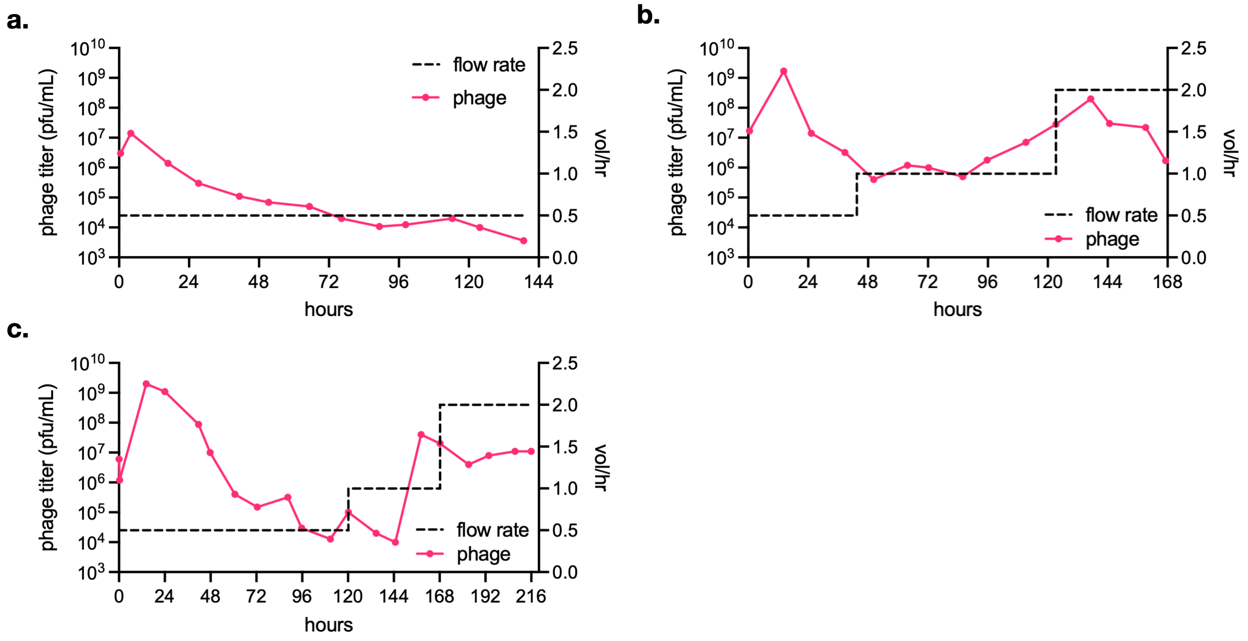


**Supplementary Figure 15.** PACE of TyrS4 phage for the formation of 3ClY, 3BrY, or 3IY. Lagoons supplemented with **a.** 500 $\mu$M 2ClPh, **b.** 200 $\mu$M 2BrPh, or **c.** 500 $\mu$M 2IPh were infected with PANCE-evolved TyrS4-phage pools (**SI Figure 14**).

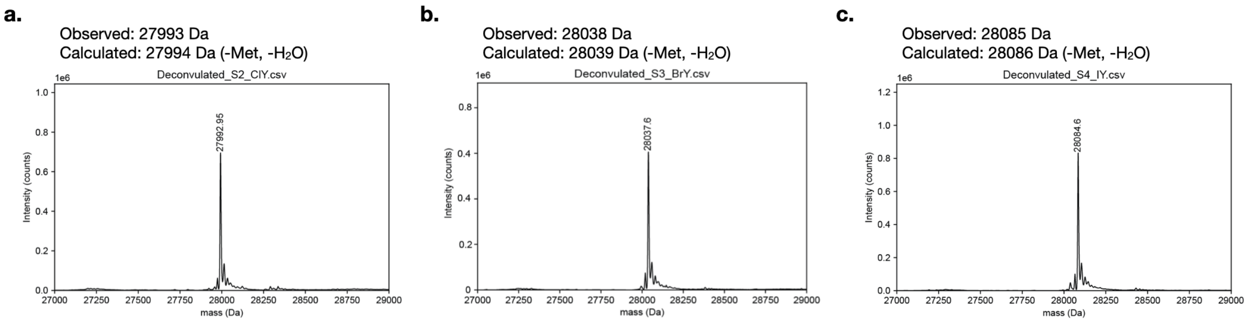


**Supplementary Figure 16.** Mass spectrometry analysis of GFP* produced in the presence of P1-1 and 500 $\mu$M **a.** 2ClPh, **b.** 2BrPh, or **c.** 2IPh. Observed masses are consistent with incorporation of the intended 3XY along with removal of the *N*-terminal Met and the loss of a water molecule from fluorophore maturation.

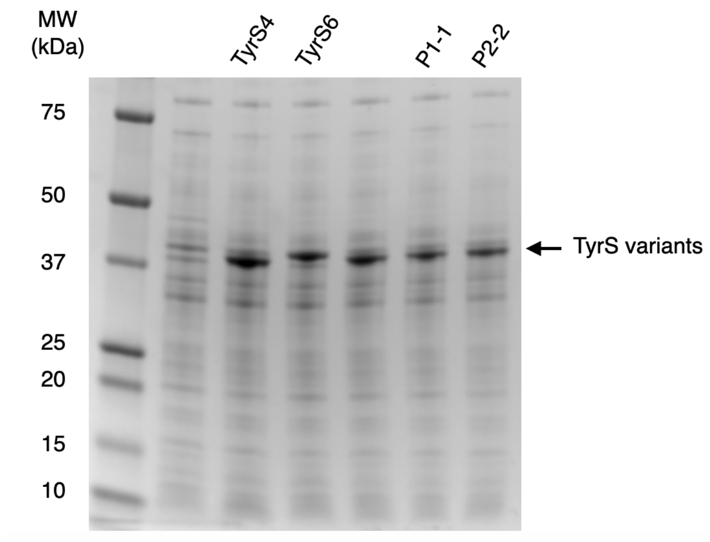


**Supplementary Figure 17.** Expression of evolved variants P1-1 and P2-1 in comparison to the parent TyrS4 and TyrS6, assessed by SDS-PAGE.

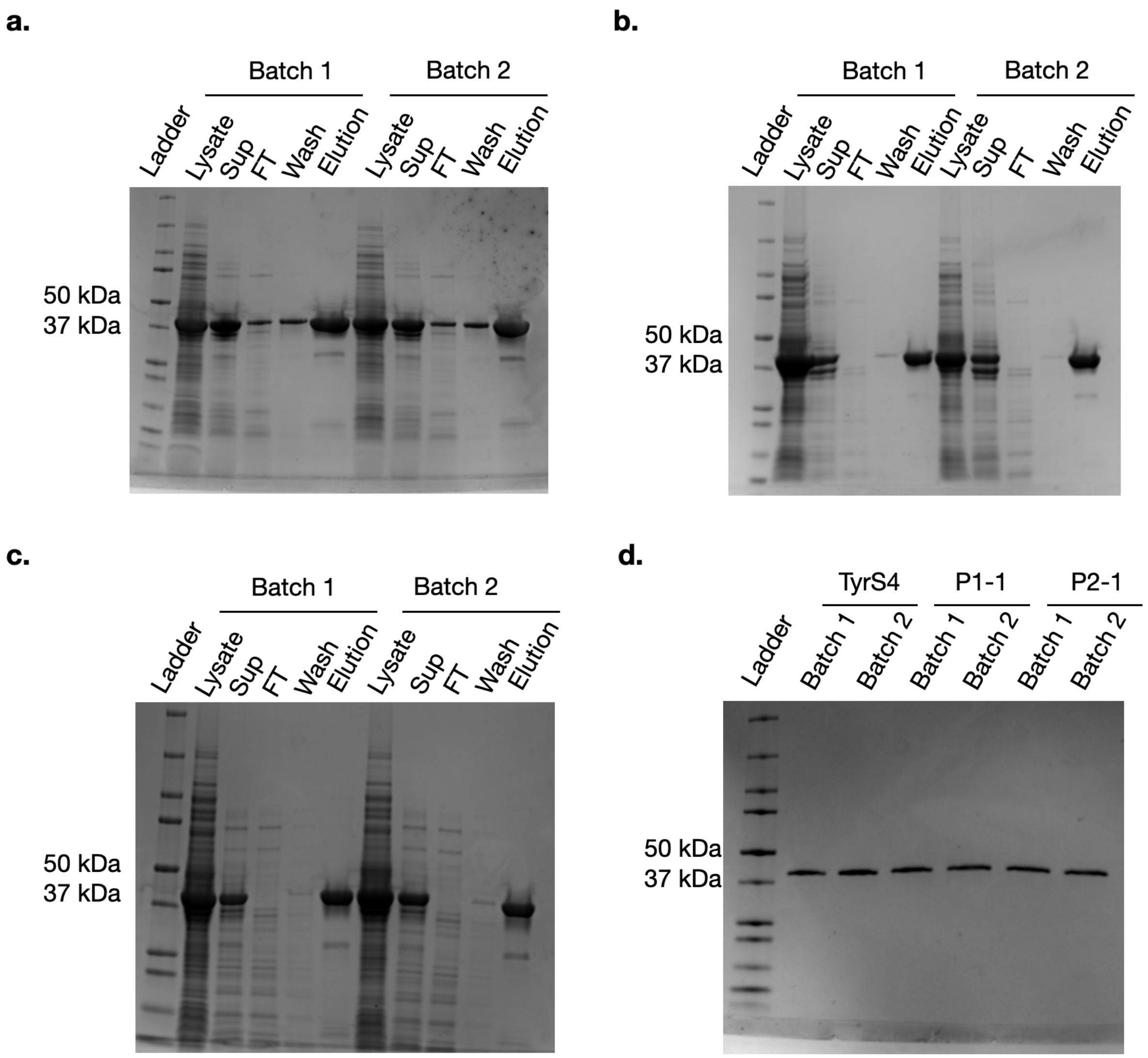


**Supplementary Figure 18.** Purification of TyrS variants. **a.-c.** SDS-PAGE gels of protein fractions from **a.** TyrS4, **b.** P1-1, and **c.** P2-1 purification. Sup = supernatant; FT = flow through. **d.** SDS-PAGE of purified enzymes.

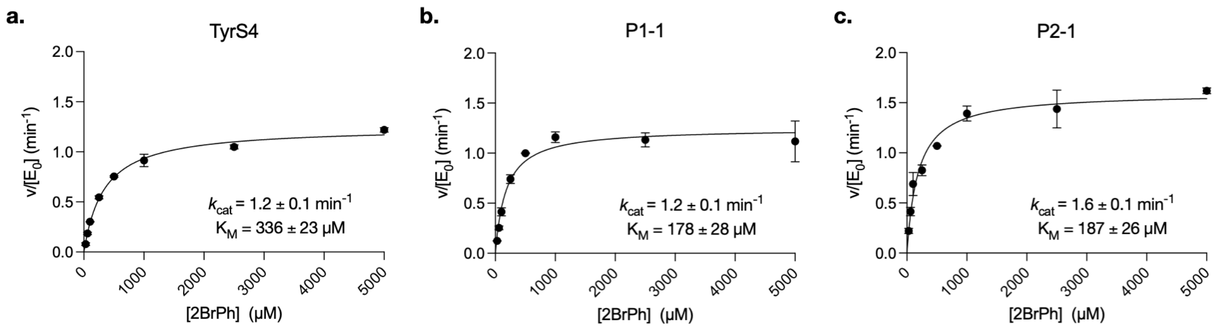


**Supplementary Figure 19.** Kinetics of TyrS4 variants for production of 3-bromotyrosine (3BrY) from 2-bromophenol (2BrPh). **a.-c.** Michaelis-Menten analysis of **a.** TyrS4, **b.** P1-1, and **c.** P2-1. Assay conditions: 1.5 $\mu$M enzyme, 5 mM L-Serine, 1 mM PLP, and variable concentrations of 2BrPh in 50 mM potassium phosphate buffer (pH 8.0) for 3 hours.

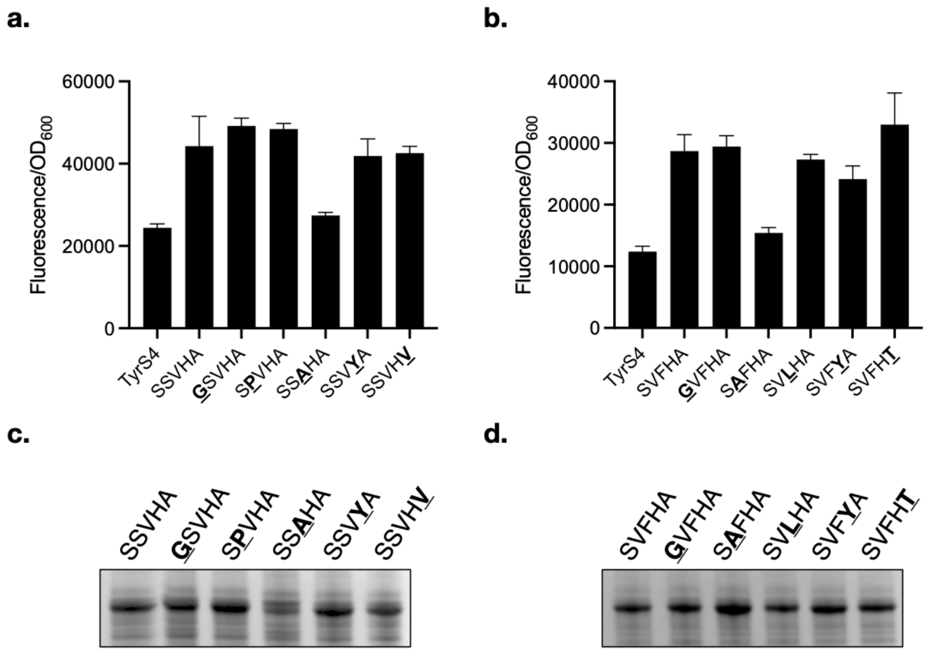


**Supplementary Figure 20. Effect of mutations in P1-1 and P2-1. a.-b.** Effect of single mutation reversions on the activities of **a.** P1-1 and **b.** P2-1 on 500 $\mu$M 2BrPh or 2IPh, respectively, through product incorporation into GFP*. **c.-d.** Assessment of single revertant expression levels by SDS-PAGE.

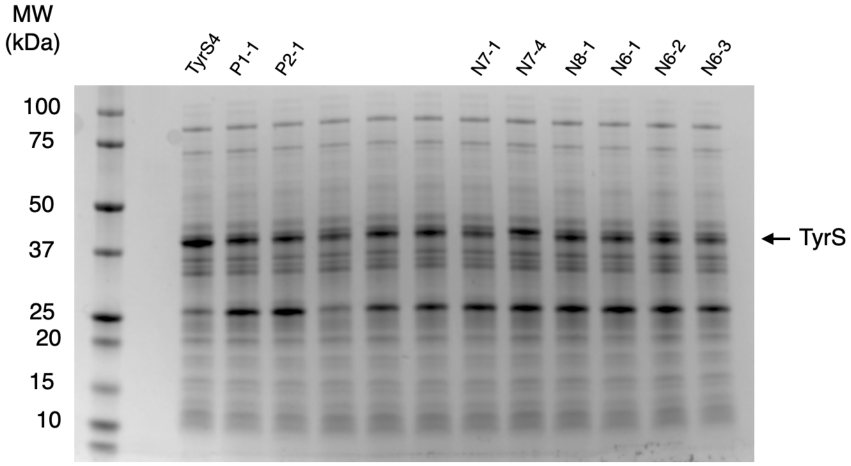


**Supplementary Figure 21.** Expression levels of TyrS4 variants isolated from the multi-site saturation library selection compared to the parent TyrS4, assessed by SDS-PAGE.

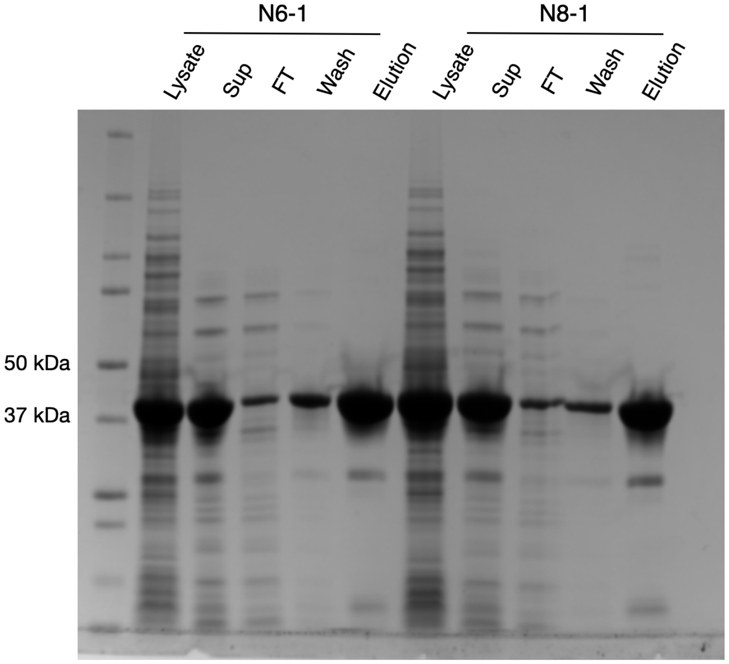


**Supplementary Figure 22.** SDS-PAGE of protein fractions from the purification of TyrS variants N6-1 and N8-1. Sup = supernatant; FT = flow through.

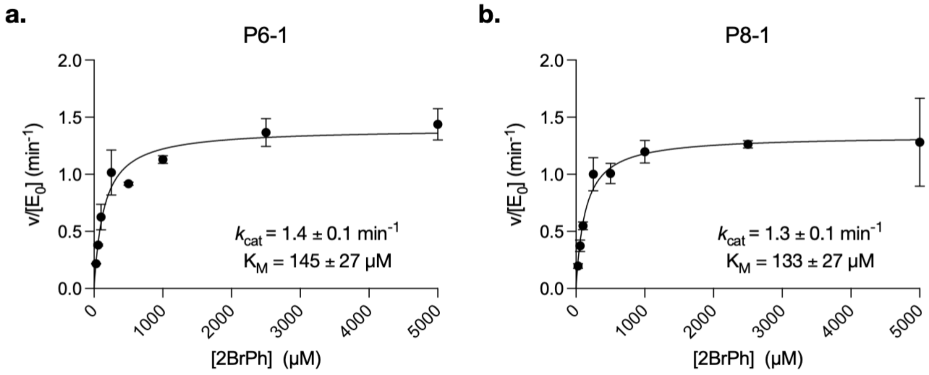


**Supplementary Figure 23.** Kinetics of TyrS4-NNK variants for production of 3-bromotyrosine (3BrY) from 2-bromophenol (2BrPh). **a.-b.** Michaelis-Menten analysis of **a.** P6-1 and **b.** P8-1. Assay conditions: 1.5 *μ*M enzyme, 5 mM L-Serine, 1 mM PLP, and variable concentrations of 2BrPh in 50 mM potassium phosphate buffer (pH 8.0) for 3 hours.

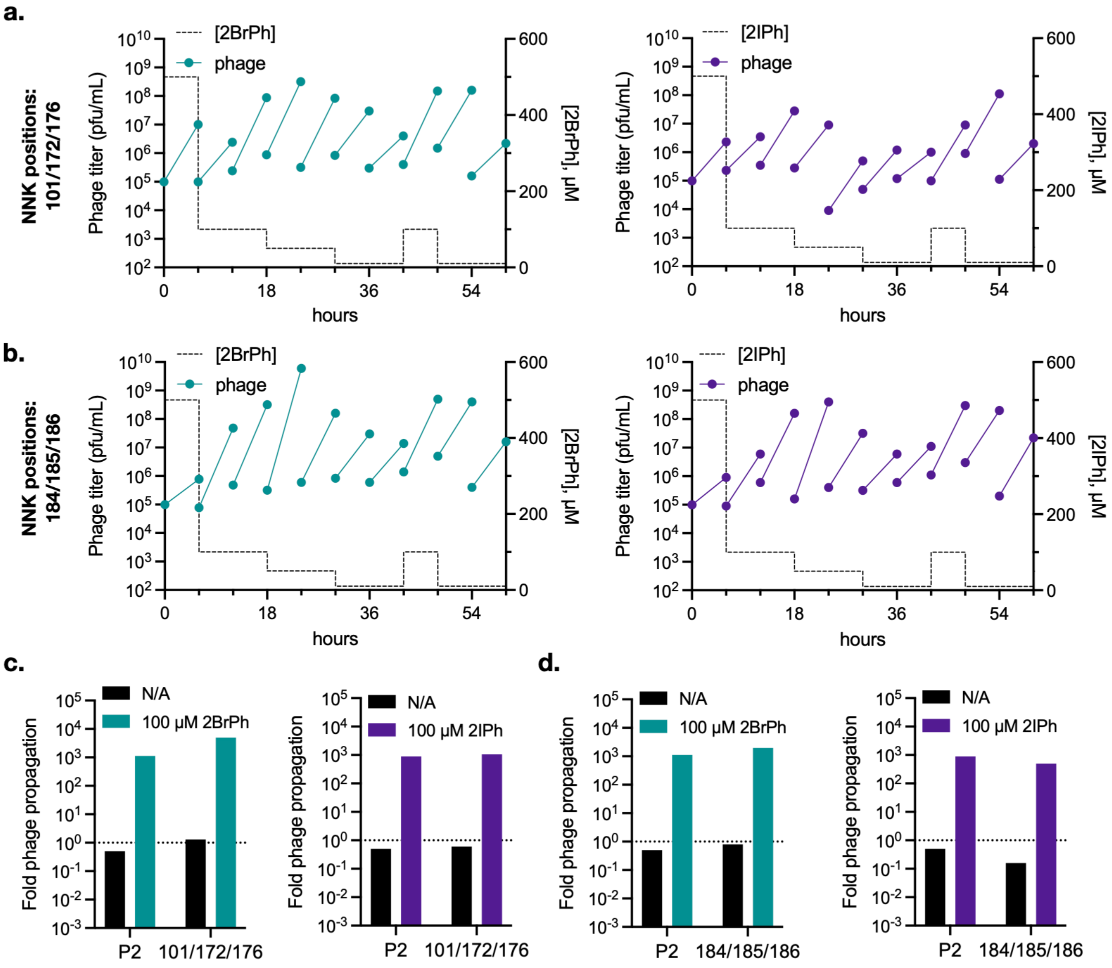


**Supplementary Figure 24.** Selection and evaluation of site-saturation mutagenesis libraries generated using PACE-evolved variant P2-1. **a.** PANCE of phage encoding a P2-1 site-saturation mutagenesis library at residues 101, 172, and 176. Phage were selected with varying concentrations of either 2BrPh (left) or 2IPh (right). **b.** PANCE of phage encoding a P2-1 site-saturation mutagenesis library at residues 184-186. **c.** Evaluation of evolved phage pools from **a**. Phage were propagated through selective host cells with no substrate ("N/A") and either 100 $\mu$M 2BrPh (left) or 2IPh (right). **d.** Evaluation of evolved phage pools from **b**. Phage were propagated through selective host cells with no substrate ("N/A") and either 100 $\mu$M 2BrPh (left) or 2IPh (right).

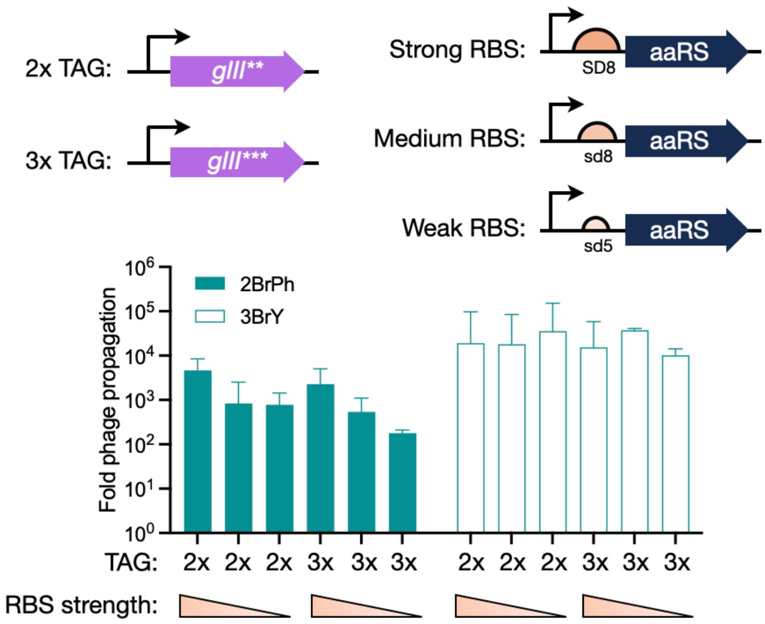


**Supplementary Figure 25.** Altering the number of TAG codons in *gIII* and the aaRS ribosome binding site (RBS) strength increases selection stringency. Top: scheme of alterations made to the selection genetic circuitry. Prior evolution experiments in this work utilized *gIII*** and the SD8 RBS to drive aaRS translation. An additional TAG stop codon within *gIII* (*gIII****), as well as medium (sd8) and weak (sd5) aaRS ribosome binding sites were examined.[1] Bottom: propagation of phage encoding PACE-evolved TyrS4 variant P2-1 under various selection conditions. Cultures were infected with phage and supplemented with either 500 $\mu$M 2BrPh or 3BrY.

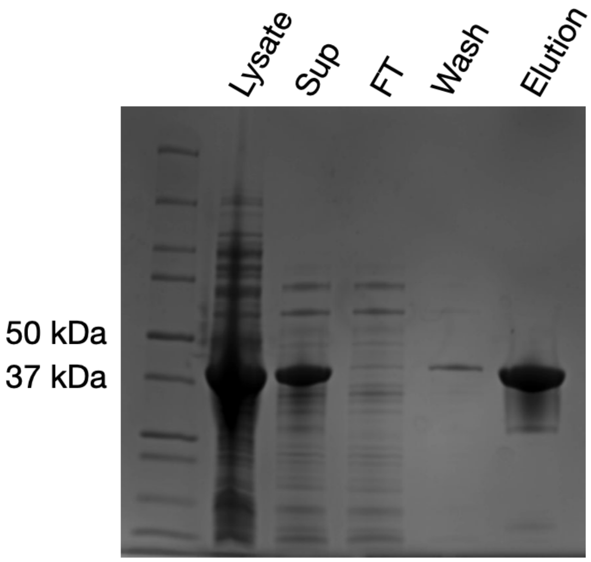


**Supplementary Figure 26.** SDS-PAGE of protein fractions from the purification of TyrS variant P3-3. Sup = supernatant; FT = flow through.

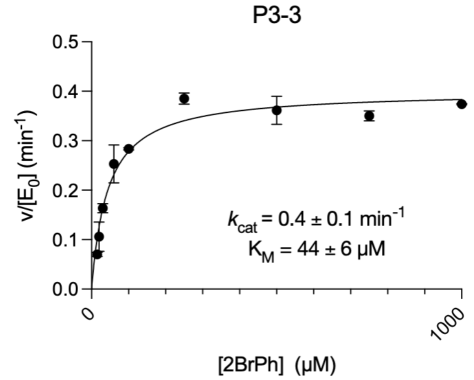


**Supplementary Figure 27.** Michaelis-Menten analysis of P3-3. Assay conditions: 1.5 $\mu$M enzyme, 5 mM L-Serine, 1 mM PLP, and variable concentrations of 2BrPh in 50 mM potassium phosphate buffer (pH 8.0) for 3 hours.
